# BMP–Smad1/9 signaling is required for PGC proliferation in zebrafish

**DOI:** 10.1101/2025.11.04.686672

**Authors:** Tao Zheng, Yaqi Li, Guangyuan Li, Zihang Wei, Jie Li, Zheng Jiang, Roshan Shah, Weiying Zhang, Cencan Xing, Anming Meng, Xiaotong Wu

## Abstract

The specification and maintenance of germ cell lineage are vital for reproductive development and heredity. The germ cell fate in some species, such as zebrafish and *Xenopus*, is determined by maternally supplied germ plasm, whereas in mammals it is induced by bone morphogenetic protein (BMP) signaling after implantation. It remains elusive whether BMP signaling is implicated in zebrafish germ cell development. Here, we demonstrate that, in zebrafish, BMP–Smad1/9 signaling is required for primordial germ cell (PGC) maintenance but not for PGC fate determination or migration. BMP inhibitor treatment or whole-embryo *smad1/9* knockdown reduces the number of PGCs. To investigate cell-autonomous roles of BMP–Smad1/9 in PGCs, we generated PGC-specific *smad1*/*9* knockouts using a double-transgenic approach with PGC-specific expression of Cas9 and ubiquitous expression of guide RNAs (gRNAs). Loss of Smad1/9 in PGCs leads to impaired PGC proliferation and increased apoptosis, resulting in reduced PGC numbers and adult sex bias. Moreover, transcriptome analysis of *smad1*-deficient PGCs revealed no significant changes in the expression of PGC-specific genes, but a marked upregulation of genes involved in cell cycle checkpoints and DNA damage repair. Notably, ectopic ATR–pChk1 activation in *smad1*-cKO PGCs validates cell cycle defects and DNA replication stress. ATR inhibition restores PGC numbers in mutant embryos. Collectively, these findings underscore a critical role of BMP–Smad1/9 signaling in zebrafish PGC population maintenance, while it is dispensable for PGC fate determination and migration.

## Introduction

The formation and transmission of germ cells are crucial for species survival and evolutionary continuity. In mammals, PGCs are induced from a subset of proximal epiblast cells around the onset of gastrulation by BMP signals from the adjacent extraembryonic ectoderm and visceral endoderm (Lawson et al., 1999; Matzuk and Burns, 2012; Cantú and Laird, 2013). Disruption of BMP signaling components, including *Bmp2*, *Bmp4*, *Bmp8b*, *Smad1*, *Smad4*, and *Smad5*, results in the complete loss or significant reduction of PGCs (Lawson et al., 1999; Ying et al., 2000; Chang and Matzuk, 2001; Ying and Zhao, 2001; Hayashi et al., 2002; Senft et al., 2019). Notably, BMP signaling has also been implicated in PGC specification in other animals, such as the hemichordate *Ptychodera flava* and the basal insect *Gryllus bimaculatus*, suggesting that BMP-mediated PGC induction may represent an evolutionarily conserved mechanism among metazoans (Donoughe et al., 2014; Lochab and Extavour, 2017; Lin et al., 2021). In contrast, PGC fate in some non-mammals, including *Caenorhabditis elegans*, *Drosophila melanogaster*, *Xenopus laevis*, and *Danio rerio*, is pre-determined by maternally inherited germ plasm containing mRNAs and proteins, such as *vasa*, *ziwi*, *dazl* (Yoon et al., 1997; Houwing et al., 2007; Jamieson-Lucy and Mullins, 2019; Bertho et al., 2021). Germ plasm is inherited and enriched in a group of cells destined for PGC fate in early embryos. After specification, PGCs migrate toward the gonadal region while undergoing proliferation (Raz, 2003). However, whether BMP signaling is essential for zebrafish PGC development remains unexplored.

The BMP signaling pathway is a crucial regulator of various biological processes, including development, cell differentiation, tissue homeostasis, cell apoptosis, stress responses, and disease progression (Morikawa et al., 2016; Zinski et al., 2018; Gomez-Puerto et al., 2019). Activation of BMP–Smad signaling triggers phosphorylation of downstream effectors Smad1, Smad5, and Smad9 (also known as Smad8), which form a complex with Smad4, translocate into the nucleus, and promote or repress gene transcription in a context-dependent manner (Shi and Massagué, 2003; Schmierer and Hill, 2007). BMP–Smad signaling is widely known to function as a morphogen during early zebrafish embryogenesis, promoting ventral cell fates (Kondo, 2007; Xue et al., 2014; Pomreinke et al., 2017; Jia and Meng, 2021; Madamanchi et al., 2021). Previous studies found that genetic disruptions of BMP ligands, such as *bmp2b* and *bmp7* (Kishimoto et al., 1997; Nguyen et al., 1998; Dick et al., 2000; Schmid et al., 2000), receptor *alk8* (Mintzer et al., 2001), and downstream effector *smad5* (Hild et al., 1999), result in dorsalized phenotypes. In addition, the non-canonical BMP signaling pathway, also named the Smad-independent pathway, has been reported to regulate some developmental processes through MAPK, PI3K–AKT, Rho–GTPase, or Wnt pathways (Moustakas and Heldin, 2005; Jia and Meng, 2021).

In this study, we show that BMP–Smad signaling is activated in zebrafish PGCs during early embryogenesis and plays a critical role in maintaining the PGC population. Inhibition of BMP signaling or whole-embryo knockdown of *smad1/9* reduced PGC numbers. Germline-specific knockout of *smad1* or *smad9* did not disrupt PGC specification or migration but impaired PGC proliferation and increased apoptosis, likely due to DNA damage response and cell cycle defects.

## Results

### BMP–Smad signaling in PGCs is required for their development

To determine whether the BMP–Smad signaling pathway plays a role in PGC development, we first examined the expression levels of BMP–Smad factors in PGCs during embryogenesis (Fig. 1A). *smad5* is highly expressed in PGCs from the high stage to 24 hours post-fertilization (hpf), while *smad1* and *smad9* show low expression before the dome stage and are then elevated (Fig. 1B). *smad4* is also expressed throughout the developmental stages while *smad1/5/9* are present before 24 hpf (Fig. 1B). Next, we detected the level of phosphorylated Smad1/5/9 (pSmad1/5/9), an activated form of Smad1/5/9, in PGCs by immunostaining. We found that pSmad1/5/9 signals were consistently detectable in PGCs, although with variable levels among individual PGCs, from the dome to 16-somite stages (Fig. 1C), indicating the existence of an active BMP–Smad signaling in PGCs during early embryogenesis.

**Fig. 1.**
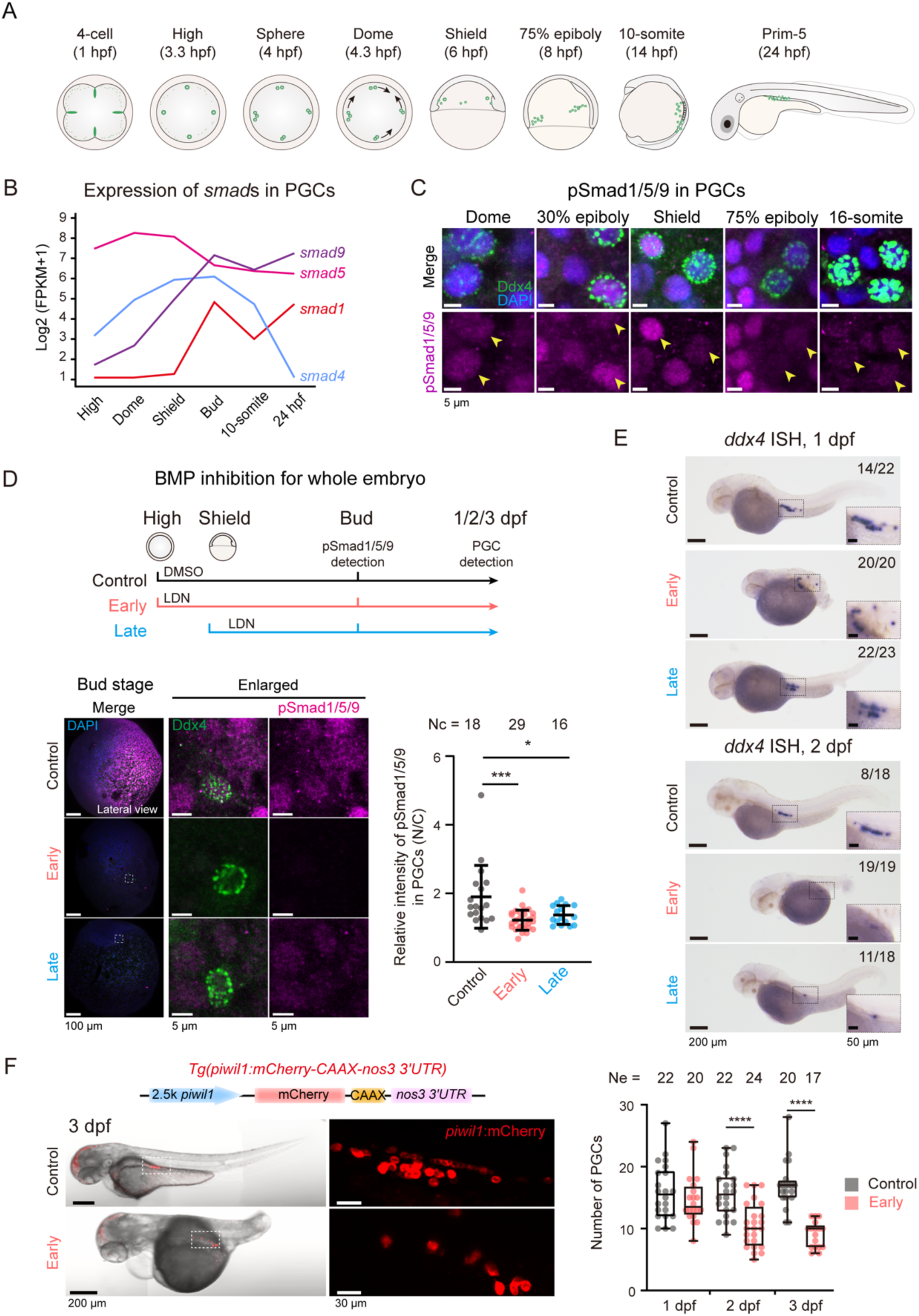
Active BMP–Smad signaling in PGCs is required for PGC development. (**A**) Schematic illustration of zebrafish PGC development. Animal pole views were shown for the 4-cell, high, sphere, and dome stages; lateral views with dorsal to the right were shown for the shield, 75% epiboly, and 10-somite stages, and with anterior to the left for the prim-5 stage. Arrows at the dome stage indicated the migration direction of PGCs. Germ plasm or germ cells were indicated in green. (**B**) Expression of related *smad* genes in PGCs during early embryogenesis (RNA-seq data from D’Orazio et al., 2021). (**C**) Immunofluorescence detection of pSmad1/5/9 (magenta) and PGC marker Ddx4 (green) at indicated stages. A selected, representative region was shown for each stage with PGCs indicated by yellow arrowheads. (**D**) Effectiveness of BMP inhibition in embryos. Upper, experimental design for BMP inhibition using the BMP inhibitor LDN-193189; lower left panel, representative images of pSmad1/5/9 detected by immunostaining; right, bar graph showing nuclear pSmad1/5/9 intensities normalized to cytoplasmic levels in individual PGCs. Each dot represents one PGC. Nc, number of observed PGCs. Error bars represented standard deviation (SD). (**E**) Visualization of PGCs by *ddx4* ISH in DMSO- or LDN-treated embryos. The ratio of embryos with representative characteristics was indicated. (**F**) Visualization and counting of PGCs in 3-dpf *Tg(piwil1:mCherry-CAAX-nos3 3’UTR)* embryos. Top left, schematic of the transgene; bottom left, representative images of embryos treated with DMSO or LDN from the high stage to 1 dpf (the “Early” group); right, box plot showing PGC numbers per embryo in each group, with the median indicated by the center line. Each dot represents one embryo. Ne, number of observed embryos. Statistical significance was determined by Student’s *t* test: *, *P* < 0.05; ***, *P* < 0.001; ****, *P* < 0.0001.

Next, to determine whether active BMP–Smad signaling is necessary for PGC development, we treated wild-type (WT) embryos with the BMP type I receptor inhibitor LDN-193189 (hereafter abbreviated as LDN), which could efficiently block phosphorylation of Smad1/5/9 (Yu et al., 2008). Given that BMP signaling is essential for dorsoventral patterning starting during blastulation (Mullins et al., 1996), we started the treatment either from the high stage (the “Early” group in Fig. 1D; Supplementary Fig. 1A) or from the shield stage (“Late” in Fig. 1D; Supplementary Fig. 1A). The latter could minimize secondary effects from disrupted patterning because dorsoventral patterning has been established at the shield stage. First of all, we assessed dorsoventral patterning in LDN-treated embryos and found that embryos in the “Early” group exhibited obvious dorsalization phenotypes, while embryos in the “Late” group showed mild dorsalization (Supplementary Fig. 1A). To further validate the BMP inhibition by LDN, we detected pSmad1/5/9 levels by immunostaining and observed a marked reduction in pSmad1/5/9 signal in both “Early” and “Late” groups at the bud stage (Fig. 1D). Then, we collected control and LDN-treated embryos at 1 and 2 days post-fertilization (dpf) for PGC detection by whole-mount *in situ* hybridization (ISH) for *ddx4* (also known as *vasa*), a germline-specific gene (Yoon et al., 1997) (Fig. 1E). The results revealed a significant reduction in PGC number in both “Early” and “Late” groups (Fig. 1E). To reliably count PGC numbers, we used *Tg(piwil1:mCherry-CAAX-nos3 3’UTR)* transgenic embryos (Li et al., 2025) for early LDN treatment, in which PGCs are specifically labeled by mCherry fluorescence driven by the 2.5-kb *piwil1* promoter and the *nanos3* (*nos3*) 3’ untranslated region (3’UTR) (Köprunner et al., 2001; Tan et al., 2002; Houwing et al., 2007; Zhang et al., 2025). Microscopic observation revealed a significant decrease in PGC numbers in LDN-treated embryos at 2 and 3 dpf (Fig. 1F). Taken together, these results suggest that active BMP signaling is essential for PGC population in zebrafish embryos.

### Knockdown of *smad1/9* results in decreased PGC number

To directly suppress BMP–Smad signaling, we chose to knock down the downstream effectors *smad1/5/9* in *Tg(piwil1:mCherry-CAAX-nos3 3’UTR)* embryos using previously validated morpholino antisense oligonucleotides (MOs) (Lele et al., 2001; Dee et al., 2007; Min et al., 2021). Triple knockdown by injecting a relatively low dose of *smad1/5/9* MOs, without causing severe embryonic dorsalization, led to a remarkable PGC reduction at 2 dpf (Fig. 2A). To determine which specific *smad* gene is responsible for PGC population maintenance, individual genes were knocked down separately. We found that *smad1* morphants, which displayed mild trunk curvature and regressed yolk stalk (Fig. 2B), as previously reported (McReynolds et al., 2007), showed a dramatic reduction in PGC number but no significant increase in mislocalized PGCs (Fig. 2B–D). In addition, immunostaining indicated that pSmad1/5/9 levels were markedly reduced in PGCs of *smad1* morphants at 75% epiboly stage (Fig. 2E). Similarly, *smad9* knockdown also led to reduced PGC number as counted at 2 dpf (Fig. 2F). In contrast, *smad5* knockdown did not affect PGC number at 2 dpf (Supplementary Fig. 1B–C). These data suggest that *smad1* and *smad9*, rather than *smad5*, are implicated in PGC population maintenance.

**Fig. 2.**
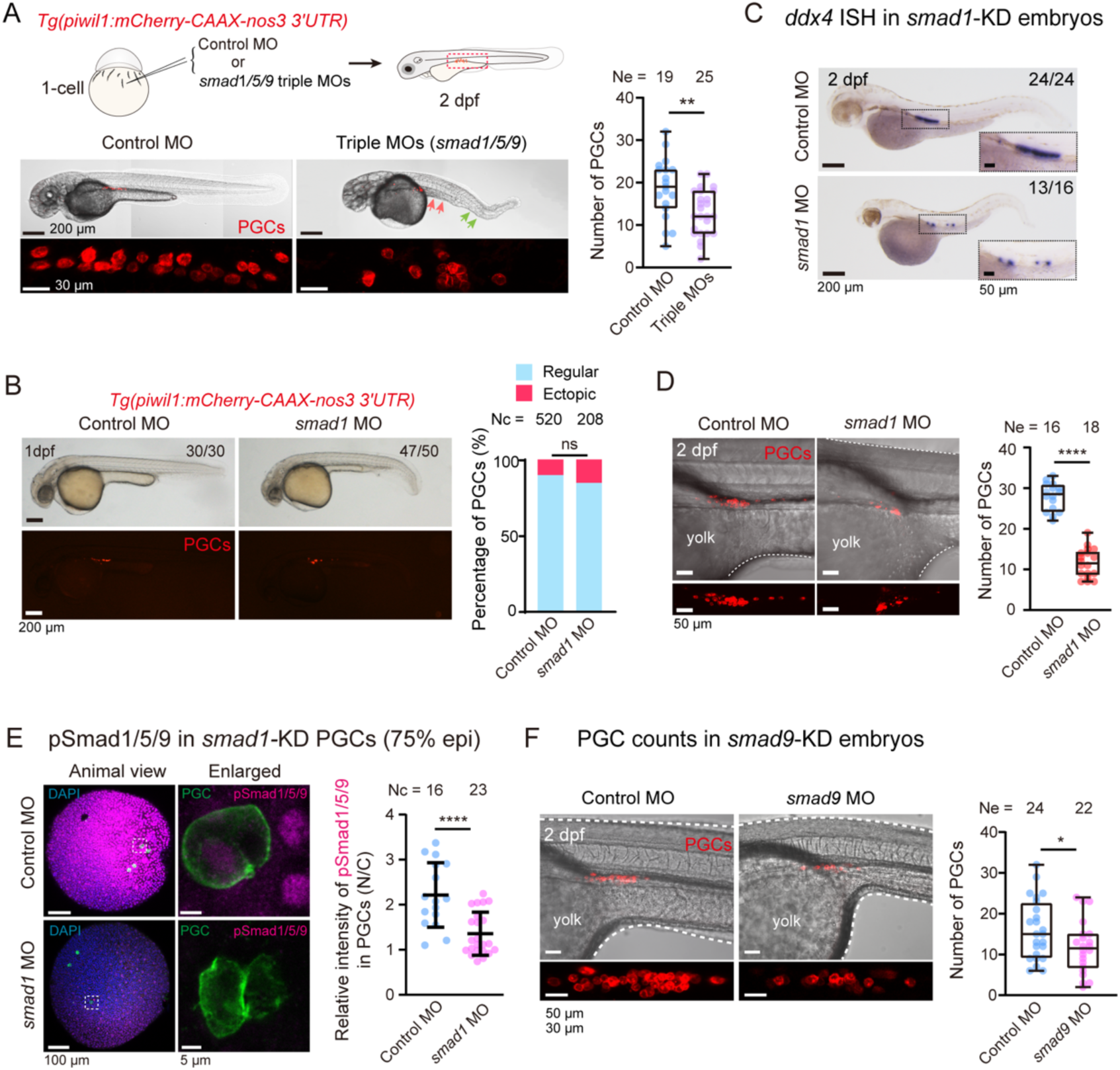
Knockdown of *smad1/5/9* leads to a reduction in PGC numbers. (**A**) Effects of *smad1/5/9* triple knockdown in *Tg(piwil1:mCherry-CAAX-nos3 3’UTR)* embryos. Injected doses were 0.5 ng *smad1* MO, 0.5 ng *smad5* MO, and 3 ng *smad9* MO. Left top, schematic of the knockdown strategy; left bottom, representative images showing embryonic morphologies and PGC numbers at 2 dpf, with red and green arrows indicating yolk stalk regression and ventral tail fin reduction, respectively; right, box plot showing PGC numbers per embryo. (**B**) Effect of *smad1* knockdown alone. Left, embryonic morphology and PGC location of representative 1-dpf embryos injected with 0.5 ng control or *smad1* MO; right, bar graph showing the percentage of mislocated PGCs. (**C**) Impact of *smad1* knockdown on *ddx4* expression in 2-dpf embryos, examined by ISH. Each embryo was injected with 0.5 ng MO at the 1-cell stage. (**D**) Impact of *smad1* knockdown on PGC numbers in *Tg(piwil1:mCherry-CAAX-nos3 3’UTR)* embryos at 2 dpf. Left, representative images with PGCs in red; right, box plot showing PGC counts per embryo. Each embryo was injected with 0.5 ng MO. (**E**) Effect of *smad1* knockdown on pSmad1/5/9 (magenta). *Tg(kop:gfp-CAAX-nos3 3’UTR)* embryos were injected with 0.5 ng MO at the 1-cell stage, and collected at 75% epiboly stage, and subjected to immunostaining with anti-GFP and anti-pSmad1/5/9 antibodies. Left, representative immunostaining images with PGCs labeled by GFP (green); right, quantification of nuclear pSmad1/5/9 intensities normalized to cytoplasmic signals, with each dot representing one PGC. Error bars represented SD. (**F**) Effect of *smad9* knockdown on PGC numbers in 2-dpf *Tg(piwil1:mCherry-CAAX-nos3 3’UTR)* embryos injected with 16 ng MO at the 1-cell stage. Left, representative images with PGCs in red; right, box plot showing PGC counts per embryo. Box-and-whisker plots displayed all values (min to max), with the median indicated by the center line. Statistical significance was determined using Student’s *t*-test unless otherwise indicated. Fisher’s exact test was used in (**B**). Statistical significance: ns, not significant (*P* ≥ 0.05); *, *P* < 0.05; **, *P* < 0.01; ****, *P* < 0.0001. Nc, number of observed PGCs. Ne, number of observed embryos.

### Generation of *smad1* and *smad9* PGC-specific mutants by *piwil1* promoter-driven *Cas9* expression

To further investigate the cell-autonomous functions of Smad1 and Smad9 in PGCs, we employed our recently developed CRISPR/Cas9-based double transgenic genome editing system that specifically induces gene mutation in germline cells (Li et al., 2025). To do this, we established two transgenic lines. The first transgenic line is *Tg(piwil1:zCas9-nos3 3’UTR)*, in which the *piwil1* promoter (Houwing et al., 2007) drives the PGC/GC-specific expression of zebrafish codon-optimized Cas9 (zCas9) (Liu et al., 2014) in the presence of *nos3* 3’UTR (Fig. 3A; Supplementary Fig. 2A). The second transgenic line ubiquitously expresses *smad1* or *smad9* gRNAs under the control of the *U6a/b/c* promoters along with *gsc:tdTomato* (tdTomato signals at dorsal region at 6 hpf) or *cryaa:CFP* (CFP signals in the lens at about 30 hpf) for easy identification of transgenic embryos (Fig. 3A; Supplementary Fig. 2B–C). Gene editing could happen in PGCs of embryos derived from crosses between these two transgenic lines (female for zCas9 and male for gRNAs), thus resulting in PGC-specific mutations of the target gene (Fig. 3A; Methods). TdTomato- or CFP-positive embryos were identified as PGC-specific KO (cKO) embryos, and negative siblings were used as controls (Fig. 3A; Supplementary Fig. 2B–C). Notably, to gain a more comprehensive observation of PGCs at different stages, we crossed *Tg(piwil1:zCas9-nos3 3’UTR)* line with *Tg(kop:Eos-CAAX-nos3 3’UTR)* or *Tg(piwil1:mCherry-CAAX-nos3 3’UTR)* for early-stage (from the dome stage to pre-2 dpf) PGC labeling or long-term (from the shield stage to adulthood) PGC labeling, respectively (Fig. 3A).

**Fig. 3.**
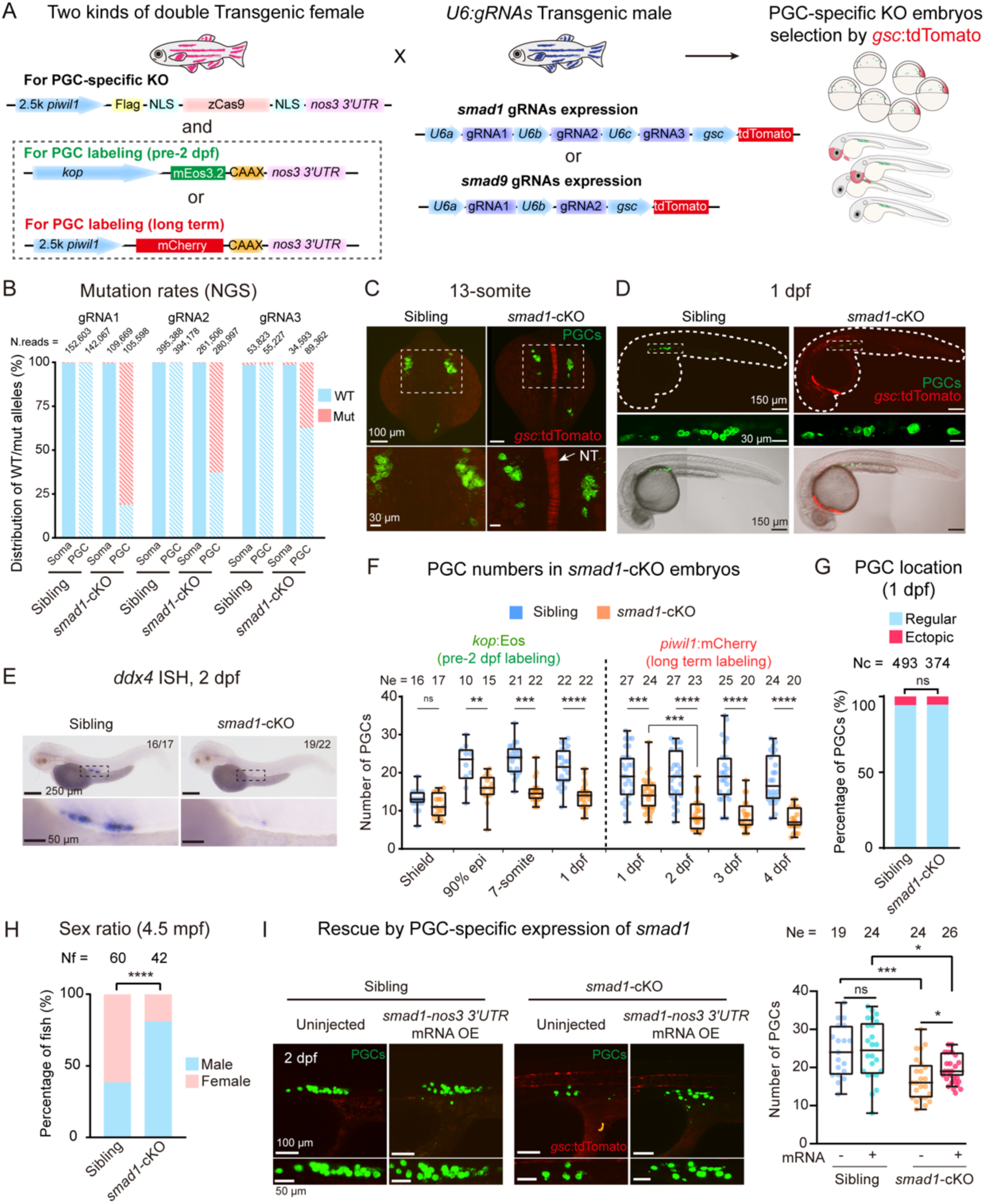
*smad1* is required cell-autonomously for PGC development. (**A**) Schematic of mating strategy used to generate *smad1*-cKO (or *smad9*-cKO) embryos. PGCs were labeled with *kop:*Eos (green fluorescence) or *piwil1:*mCherry (red fluorescence), and cKO embryos identified by *gsc:*tdTomato (in the dorsal mesoderm) or *cryaa:*CFP (in the eye lens) selection markers. (**B**) Genome editing efficiency of three *smad1* gRNA target sites assessed by targeted amplicon sequencing in somatic cells and PGCs sorted from control and *smad1*-cKO embryos at 15-somite stage. N. reads, number of aligned reads per target site. (**C** and **D**) Representative images of PGCs in control and *smad1*-cKO embryos at 13-somite stage (**C**) and 1 dpf (**D**). Embryos were dorsally viewed with anterior to the top (**C**), or laterally viewed with anterior to the left (**D**). PGCs were *kop:*Eos-positive (green). The *gsc:*tdTomato-positive (red) notochord was indicated by arrows (**C**). (**E**) ISH-detected *ddx4* expression in control and *smad1*-cKO embryos at 2 dpf. The boxed gonad region in the upper panel was enlarged in the lower panel. (**F**) Quantification of PGCs per embryo in control and *smad1*-cKO groups at the indicated stages. (**G**) The proportion of mislocated PGCs in control and *smad1*-cKO embryos at 1 dpf. (**H**) Sex ratios of control and *smad1*-cKO adult fish at 4.5 months post-fertilization (mpf). (**I**) Rescue of PGC numbers in *smad1*-cKO embryos by *smad1-nos3 3’UTR* mRNA injection (250 pg/embryo). Left, representative fluorescent images of 2-dpf embryos (similar to those in (**C** and **D**)) with *kop:*GFP-positive PGCs (green); right, box plot showing PGC numbers per embryo. Box-and-whisker plots (**F**, **I**) showed all values (min to max), with the median indicated by the center line. Each dot represented one embryo. Statistical analysis was performed using Student’s *t*-test (**F**, **I**) or Fisher’s exact test (**G**, **H**). Statistical significance: ns, not significant (*P* ≥ 0.05); *, *P* < 0.05; **, *P* < 0.01; ***, *P* < 0.001; ****, *P* < 0.0001. Ne, number of embryos analyzed; Nc, number of PGCs quantified; Nf, number of fish tested.

To validate the genome editing efficiency of our PGC-specific KO system, we first utilized FACS to sort mCherry-labeled PGCs in control siblings (with zCas9 in PGCs but without gRNAs) and *smad1*-cKO embryos at 2 dpf (Supplementary Fig. 2D). Quantitative RT-PCR (qRT-PCR) confirmed that *ddx4* was markedly enriched in sorted PGCs relative to somatic cells (Supplementary Fig. 2D), indicating effective sorting. Next, we assessed the mutation efficiency at the target regions of *smad1* in both PGCs and somatic cells by performing whole genome amplification (WGA) on a limited number of FACS-sorted cells at the 15-somite stage, followed by PCR amplification of genome regions flanking the three *smad1* gRNA target sites and next generation sequencing (NGS). Sequencing results revealed that 81.10% (n = 85,644/105,598) reads of the gRNA1 target region were mutated in PGCs of *smad1-*cKO embryos, whereas only 0.26% (n = 397/152,603), 0.31% (n = 435/142,067), and 0.48% (n = 522/109,669) of reads were mutated in sibling somatic cells, sibling PGCs, and *smad1*-cKO somatic cells, respectively (Fig. 3B; Supplementary Fig. 2E). For the gRNA2 target region, a relatively high mutation rate (62.65%, n = 176,032/280,997) was found in *smad1*-cKO PGCs, compared with 0.16% (n = 409/261,506) in *smad1*-cKO somatic cells, 0.14% (n = 553/395,388) in sibling somatic cells, and 0.16% (n = 616/394,178) in sibling PGCs (Fig. 3B; Supplementary Fig. 2E). For the gRNA3 target region, we observed a moderate editing rate in *smad1*-cKO PGCs (37.22%, n = 33,258/89,362), and extremely low mutation frequencies in *smad1*-cKO somatic cells (1.38%, n = 478/34,593), sibling somatic cells (1.39%, n = 747/53,823), and sibling PGCs (1.28%, n = 707/55,227) (Fig. 3B; Supplementary Fig. 2E). Together, these results prove that PGC-specific knockout system works effectively. It is worth mentioning that, as previously reported (Li et al., 2025), biallelic loss-of-function mutations are likely present in only a fraction of PGCs.

### PGC-specific knockout of *smad1* and/or *smad9* leads to reduced PGC numbers

Next, we examined PGC development in *smad1-* or *smad9*-cKO embryos with PGCs labeled by *kop:*Eos expression at pre-2 dpf or by *piwil1:*mCherry expression after 2 dpf (Fig. 3A). First, we counted the number of PGCs in *smad1*-cKO embryos from the shield stage to 4 dpf and observed a significant lower PGC numbers in *smad1*-cKO embryos compared to siblings from the 90% epiboly stage through 4 dpf (Fig. 3C–F). The magnitude of the decline in PGC numbers ranged from 27.77% to 57.35% (Fig. 3F), which is reasonable since only a subset of PGCs is expected to carry biallelic null mutations through our transient editing system. Despite reduced PGC numbers, overall embryonic morphology, including axis formation, remained unaffected in *smad1*-cKO embryos at 1 and 2 dpf (Fig. 3D–E; Supplementary Fig. 2B), consistent with the extremely low mutagenesis rates observed in somatic cells of *smad1*-cKO embryos (Fig. 3B; Supplementary Fig. 2E). Notably, the majority of *smad1*-cKO PGCs migrated normally to the future gonads at 1 dpf (Fig. 3E, G). Given that PGC numbers influence sex determination in zebrafish with a higher number of PGCs promoting female development and a lower number skewing towards males (Tzung et al., 2014; Dranow et al., 2016; Ye et al., 2019), we raised these *smad1*-cKO embryos and their siblings to adulthood (4.5 months post-fertilization) and found that *smad1*-cKO fish exhibited a strong male bias in sex determination (Fig. 3H). Importantly, overexpression of synthetic *smad1-nos3 3’UTR* mRNA, which could be specifically translated in PGCs, partially recovered PGC numbers in *smad1*-cKO embryos (Fig. 3I). Therefore, these data suggest that loss of Smad1 in PGCs would trigger a decline in PGC numbers.

Similarly, *smad9*-cKO embryos showed a 39.68% decrease in PGC number at 3 dpf (Fig. 4A–B). In addition, *smad1&9* double-cKO embryos exhibited an even greater decrease in PGC number compared to either *smad1*-cKO or *smad9*-cKO embryos as counted at 2 dpf (Fig. 4C). Thus, we concluded that Smad1 and Smad9 in PGCs are critical for forming normal PGC numbers in zebrafish.

**Fig. 4.**
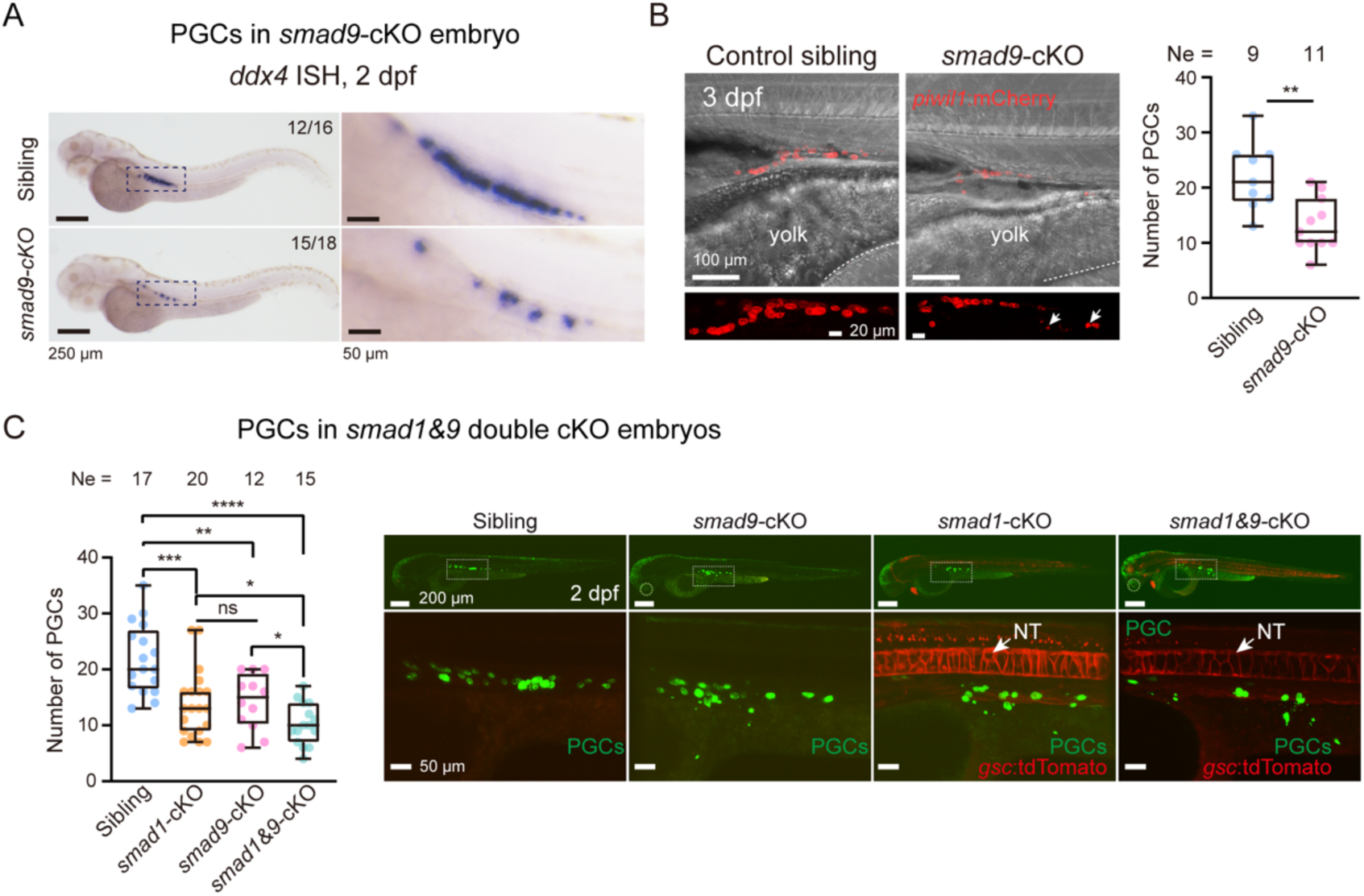
PGC-specific knockout of *smad9* leads to reduced PGC numbers. (**A**) Representative *ddx4* ISH images of control and *smad9*-cKO embryos. The boxed gonad region in the left panel was enlarged in the right panel. (**B**) Effect of *smad9*-cKO on PGC number. Left, representative images showing PGC development in 3-dpf control and *smad9*-cKO embryos. White arrows indicated cellular debris. Right, box plot showing PGC numbers per embryo. (**C**) Quantification of PGC numbers in different mutants. Left, box plot showing PGC numbers per embryo in control siblings, *smad1*-cKO, *smad9*-cKO, and *smad1&9*-cKO embryos at 2 dpf. Right, representative images of PGCs in the corresponding embryos. PGCs were labeled by *kop:*Eos. Dotted circles marked *cryaa:*CFP fluorescence in the lens of *smad9*-cKO embryos; red fluorescence indicated *gsc:*tdTomato in *smad1*-cKO embryos. NT, notochord. Box-and-whisker plots (**B**, **C**) showed all values (min to max), with the median indicated by the center line. Each dot represented one embryo. Statistical analysis was performed using Student’s *t*-test. Statistical significance: ns, not significant (*P* ≥ 0.05); *, *P* < 0.05; **, *P* < 0.01; ***, *P* < 0.001; ****, *P* < 0.0001. Ne, number of embryos analyzed.

### Deficient PGC proliferation and ectopic apoptosis caused by PGC-specific loss of Smad1

To determine whether reduced PGC numbers in PGC-specific *smad1/9* KO mutants result from impaired cell proliferation and/or apoptosis, we traced *kop:*Eos-labeled PGCs via live imaging within *smad1*-cKO embryos and siblings from 50% to 90% epiboly stages (Fig. 5A). From the germ ring to 90% epiboly stages, 51.43% (n = 18/35) of imaged PGCs in siblings exhibited cell division, while only 10.53% (n = 2/19) of the imaged PGCs in *smad1*-cKO embryos underwent cell division (Fig. 5B). To validate defective PGC proliferation, we performed EdU incorporation in *smad1*-cKO embryos and siblings from the shield to bud stages and collected embryos at 2-somite stage for observation of *kop:*Eos-labeled PGCs (Fig. 5C). The majority (81.91%, n = 77/94) of PGCs in siblings showed EdU signals, indicating active DNA replication and cell proliferation (Fig. 5C). In contrast, significantly fewer (62.50%, n = 45/72) *smad1*-cKO PGCs exhibited EdU signals, suggesting deficient cell proliferation of *smad1*-cKO PGCs (Fig. 5C). The higher proliferation rate detected by EdU compared to live imaging (Fig. 5B–C) likely reflects the greater sensitivity of EdU to DNA synthesis. Additionally, live imaging captures fewer cells for analysis due to cell migration and embryo depth. We observed numerous cellular fragments with *piwil1:*mCherry fluorescence in *piwil1:mCherry;smad1*-cKO embryos at 2 dpf (Fig. 5D, asterisk) and *piwil1:mCherry;smad9*-cKO embryos (Fig. 4B, arrow). Live imaging of *piwil1:mCherry*;*smad1*-cKO PGCs captured the process in which PGCs shed abundant debris (Fig. 5D). Via immunostaining of active Caspase-3, PGC apoptosis was evidenced by the increased active Caspase-3 signals in *smad1-*cKO PGCs, compared to siblings (Fig. 5E). Therefore, these results indicate that impaired PGC proliferation and increased apoptosis of PGCs most likely account for the reduced PGC numbers in *smad1*-cKO embryos.

**Fig. 5.**
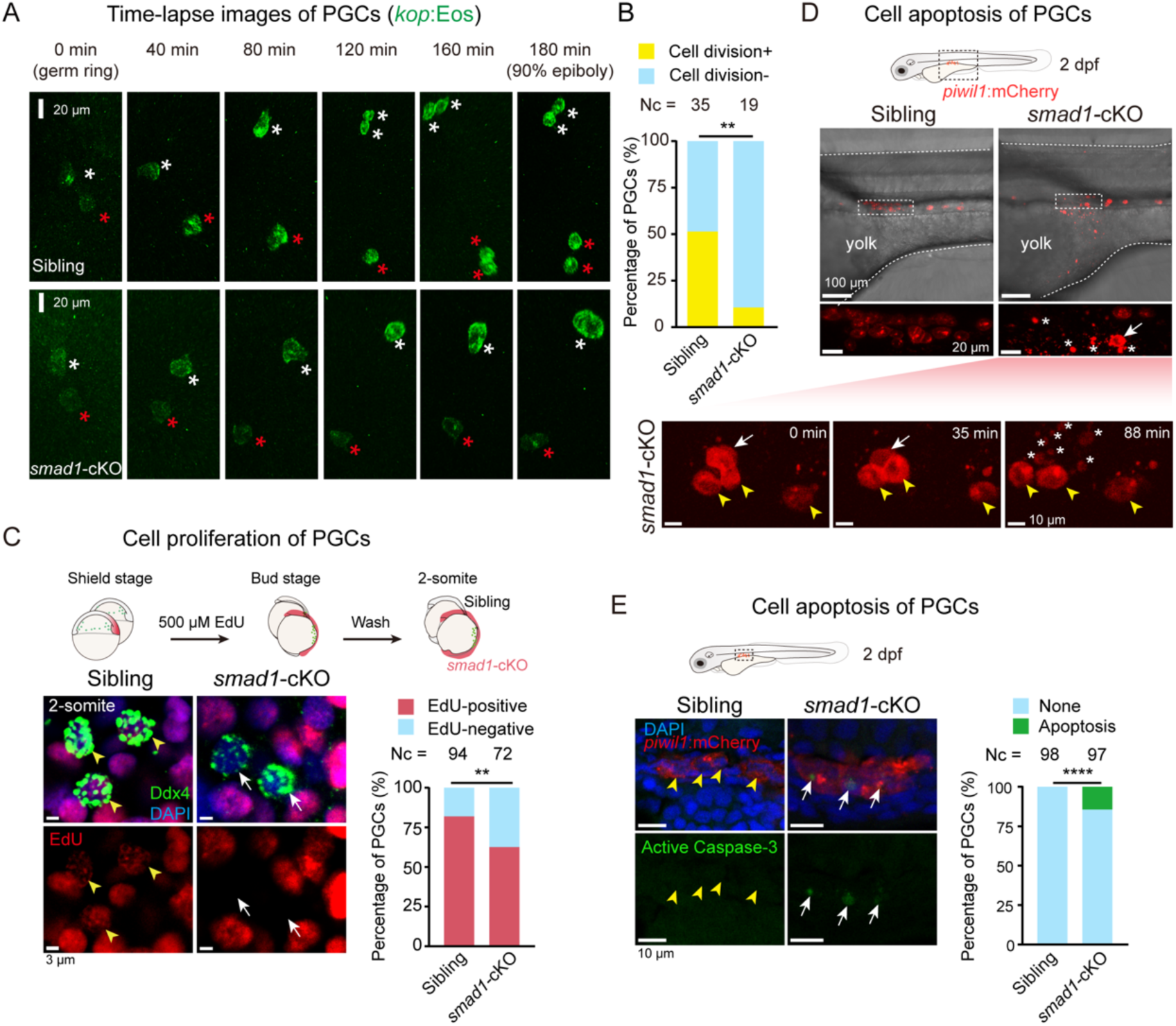
*smad1*-cKO PGCs exhibit reduced proliferation and increased apoptosis. **(A)** Representative tracking images of dividing PGCs in control and *smad1*-cKO embryos from germ ring to 90% epiboly stage. The *kop:*Eos-labeled PGCs were marked by red and white asterisks. **(B)** Ratio of dividing PGCs in embryos shown in (**A**). (**C**) Detection of proliferating PGCs via EdU incorporation assay. Embryos at shield stage were treated with EdU until the bud stage and collected at 2-somite stage for immunostaining with EdU and Ddx4 antibodies. Top panel, schematic of assay procedure; lower left, representative images of PGCs. Ddx4- and EdU-positive PGCs were indicated by yellow arrowheads, and Ddx4-positive but EdU-negative PGCs by white arrows. Lower right, bar graph showing the percentage of EdU-positive and EdU-negative PGCs in control and *smad1*-cKO embryos. (**D**) Live images of *smad1-*cKO PGCs undergoing apoptosis. Top panel, *piwil1:*mCherry-labeled PGCs in the gonad region; bottom panel, time-lapse images showing PGCs undergoing apoptosis. Yellow arrowheads marked surviving PGCs, white arrows indicated apoptotic PGCs, and asterisks denoted apoptotic cell debris. (**E**) Detection of apoptotic PGCs by immunostaining for active Caspase-3. Left panel, representative images of PGCs in the gonad region of control and *smad1*-cKO embryos at 2 dpf. PGCs were labeled by *piwil1:*mCherry. Active Caspase-3–positive PGCs were indicated by white arrows, and active Caspase-3–negative PGCs were indicated by yellow arrowheads. Right, bar graph showing the percentage of PGCs positive or negative for active Caspase-3 in each group. Statistical significance was determined using Fisher’s exact test: **, *P* < 0.01; ****, *P* < 0.0001. Nc, number of observed PGCs.

### Cell cycle defects and DNA damage response in *smad1* deficient PGCs

To investigate molecular mechanisms underlying BMP–Smad signaling regulation in PGC population maintenance during early embryonic development, we performed RNA-seq of Eos-positive PGCs sorted by FACS from *kop:Eos;smad1*-cKO embryos and siblings, respectively, at 75% epiboly and 1 dpf stages with three biological replicates (Fig. 6A; Supplementary Fig. 3A, left). First, we checked the expression levels of PGC-specific genes such as *nanos3*, *ddx4*, *ca15b*, and *dnd1* (Yoon et al., 1997; Beer and Draper, 2013; Hartwig et al., 2014; Gross-Thebing et al., 2017) and found no significant changes in *smad1*-cKO PGCs (Fig. 6B). Moreover, we performed RNA-seq of sorted PGCs from *smad1* MO and control MO injected embryos at 75% epiboly and 1 dpf stages with two biological replicates (Supplementary Fig. 3A, right), and found that expression levels of PGC-specific genes in *smad1*-KD PGCs were also comparable to those in control PGCs (Supplementary Fig. 3B). Collectively, these data suggest that Smad1 is unlikely to participate cell-autonomously in zebrafish PGC fate determination.

**Fig. 6.**
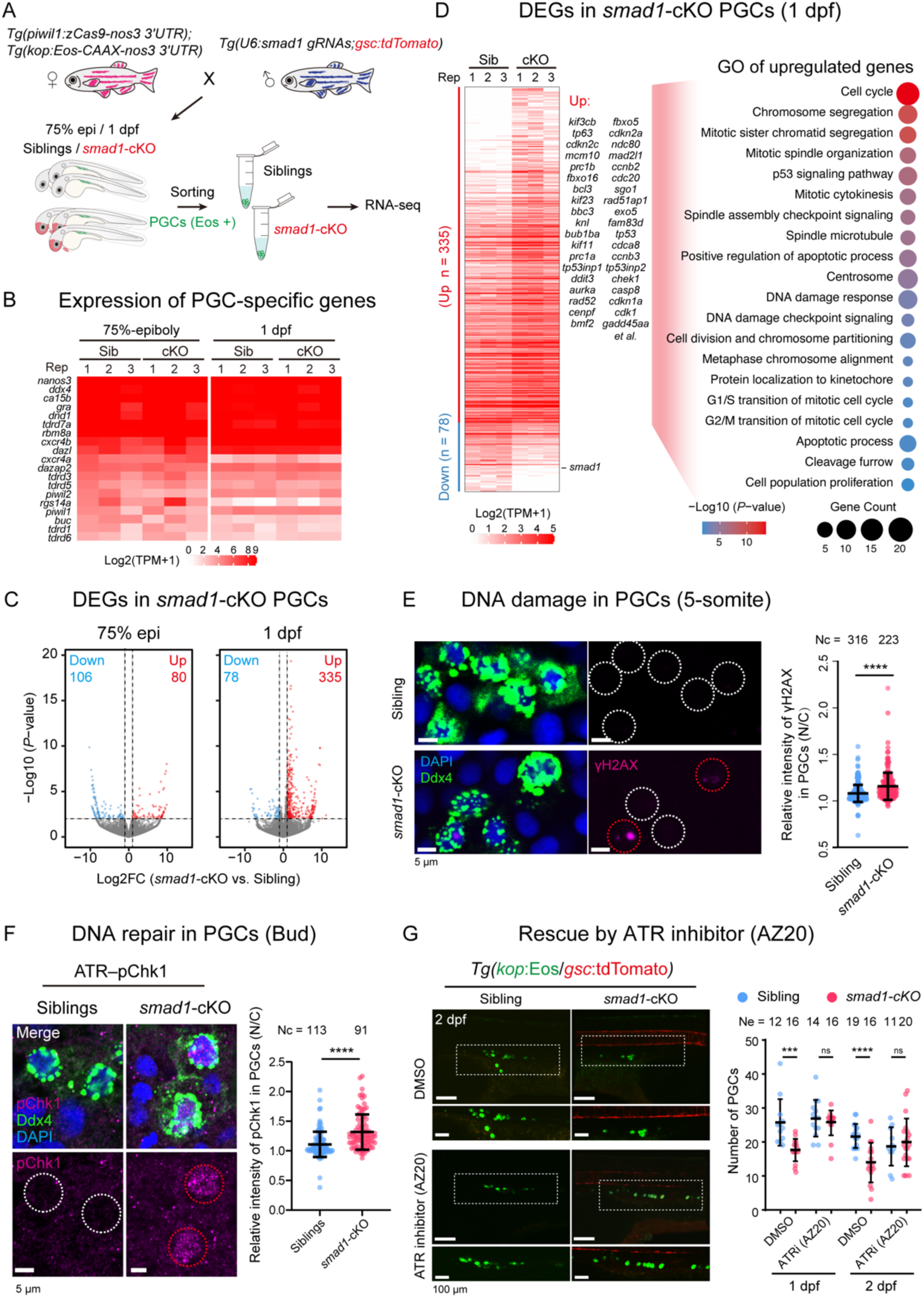
Reduced PGC number in *smad1*-cKO embryos is attributed to DDR and can be rescued by ATR pathway inhibition. (**A**) Schematic illustration of experimental design for isolation and RNA-seq of control and *smad1*-cKO PGCs. (**B**) Expression levels of PGC-specific genes in *smad1*-cKO and control PGCs based on RNA-seq data. (**C**) Volcano plots showing DEGs between *smad1*-cKO and control PGCs at indicated stages. Red and blue dots indicated upregulated and downregulated genes, respectively. (**D**) Heatmap (left) and enriched GO terms (right) of DEGs between *smad1*-cKO and control PGCs at 1 dpf. Genes related to apoptosis and cell division, along with *smad1*, were indicated. The bubble plot showed enriched GO terms for upregulated genes in *smad1*-cKO PGCs. (**E**) Detection of DNA damage in PGCs via γH2A immunostaining assay in sibling and *smad1*-cKO PGCs at the 5-somite stage. Left, representative images with PGCs labeled by Ddx4. Red and white dotted circles indicated γH2A-positive and -negative PGCs, respectively. Right, quantification of nuclear γH2A intensities normalized to cytoplasmic levels. Each dot represented one PGC. (**F**) Detection of pChk1 activation by immunostaining in control and *smad1*-cKO PGCs at the bud stage. Left, representative images with PGCs labeled by Ddx4. Red and white circles indicated pChk1-positive and -negative PGCs, respectively. Right, quantification of nuclear pChk1 intensities normalized to cytoplasmic levels. Each dot represented one PGC. (**G**) Representative images (left) and quantification of PGCs (right) in the gonad regions of control and *smad1*-cKO embryos treated with DMSO or ATR inhibitor (AZ20) from the sphere stage until 1 or 2 dpf. PGCs were labeled with *kop:*Eos. Each dot represented one embryo. Error bars represented SD. Statistical significance was determined by Student’s *t*-test: ns, not significant (*P* ≥ 0.05); ***, *P* < 0.001; ****, *P* < 0.0001. Nc, number of observed PGCs; Ne, number of observed embryos.

Next, detailed analyses identified 80 upregulated genes and 106 downregulated genes (*P* < 0.01, fold change (FC) > 2) at 75% epiboly stage and 335 upregulated genes and 78 downregulated genes at 1 dpf in *smad1-*cKO PGCs (Fig. 6C; Supplementary Table 1), indicating mild changes in the transcriptome of *smad1-*cKO PGCs at 75% epiboly stage (Fig. 6C, left). The most prominent gene expression change caused by loss of Smad1 in PGCs was the upregulation of 335 genes in cKO PGCs at 1 dpf (Fig. 6C, right). Gene ontology (GO) analysis of these 335 upregulated genes revealed that the GO terms mitotic cell cycle, chromosome segregation, mitotic checkpoint activation, and DNA damage response (DDR) were significantly enriched (Fig. 6D), which might be responsible for cell cycle defects and cell apoptosis of *smad1*-cKO PGCs. Furthermore, to validate the DNA damage in *smad1-*cKO PGCs, we conducted γH2AX immunostaining and observed elevated γH2AX signals in *smad1*-cKO PGCs (Fig. 6E). Meanwhile, to exclude non-specific DNA damage that might be caused by CRISPR/Cas9 genome editing, we also analyzed differentially expressed genes (DEGs) in *smad1*-KD PGCs, and identified 69 upregulated genes and 111 downregulated genes at 75% epiboly stage, and 394 upregulated genes and 687 downregulated genes at 1 dpf in *smad1-*KD PGCs (Supplementary Fig. 3C). However, the overlapping DEGs between cKO and KD PGCs were quite limited, and many developmental genes in KD PGCs were also affected (data not shown), which may reflect distinct consequences of global BMP signaling disruption in *smad1*-KD embryos compared to PGC-specific Smad1 deficiency in *smad1*-cKO embryos. Notably, despite limited overlaps, many genes involved in DNA damage repair and cell apoptosis, such as *tp53*, *casp8*, *sesn3*, *foxo3b*, *gadd45aa*, *gadd45ba*, *tp53inp1*, *ccng1*, and *phlda3*, were also upregulated in *smad1*-KD PGCs (Supplementary Fig. 3D; Supplementary Table 1), hinting that *smad1* deficiency might lead to excessive DDR in PGCs.

### Aberrant activation of ATR was responsible for *smad1*-cKO PGC reduction

Ataxia telangiectasia and Rad3-related (ATR) is the apical DNA replication stress response kinase, specifically in proliferating cells (de Klein et al., 2000; Blackford and Jackson, 2017). A key role of ATR is to phosphorylate and activate the protein kinase Chk1 (Guo et al., 2000; Liu et al., 2000), slowing or arresting cell cycle progression when DNA damage occurs, which then allows more time for DNA repair to prevent premature mitotic entry (Brown and Baltimore, 2003; Patil et al., 2013). However, if the DNA damage is too extensive, ATR–Chk1 activation could directly lead to cell apoptosis (Roos et al., 2016; Liu et al., 2018). To determine whether ATR–Chk1 is aberrantly activated in *smad1*-cKO PGCs in response to DNA damage, we assessed phosphorylated Chk1 (pChk1) level via immunostaining and found that *smad1*-cKO PGCs exhibited significantly elevated pChk1 signals (Fig. 6F). Notably, we attempted to rescue PGC apoptosis using AZ20, an ATR inhibitor that has been shown to restore erythrocyte number by counteracting DNA damage-induced cell cycle arrest and apoptosis in zebrafish erythroid progenitors (Weinreb et al., 2022). Interestingly, AZ20 treatment could restore PGC numbers in *smad1*-cKO embryos at both 1 and 2 dpf (Fig. 6G, red). AZ20-treated siblings showed no obvious changes in PGC numbers (Fig. 6G, blue). Thus, these data indicate that ectopic activation of ATR–Chk1 in response to DNA damage might trigger cell proliferation arrest and apoptosis of *smad1*-cKO PGCs.

In addition, we tested whether another major DDR pathway is involved—the ataxia-telangiectasia mutated (ATM)–Chk2 pathway, which primarily responds to DNA double-strand breaks. To this end, we assessed the level of phosphorylated Chk2 (pChk2), a marker for ATM activation (Maréchal and Zou, 2013), in mutant PGCs. The pChk2 signals were hardly detected in PGCs of either *smad1*-cKO mutants or siblings (Supplementary Fig. 3E). Furthermore, treatment with KU60019, an ATM inhibitor, could not restore PGC numbers in *smad1*-cKO mutants (Supplementary Fig. 3F), underscoring that ATM–Chk2 signaling may not be involved in PGC reduction in *smad1*-cKO mutants.

### Alternative splicing of genes involved in DNA damage response and cell cycle in *smad1*-deficient PGCs

There were only minor changes in the transcription levels due to the loss of Smad1 in PGCs at 75% epiboly stage (Fig. 6C). Notably, increasing evidence has shown that Smads, beyond their functions in transcriptional regulation, also participate in post-transcriptional regulation across multiple species (Davis et al., 2008, 2010; Tripathi et al., 2016). Therefore, we investigated whether Smad1 contributes to post-transcriptional regulation by performing transcriptome-wide analysis of alternative splicing (AS) in *smad1*-cKO PGCs using replicate multivariate analysis of transcript splicing (rMATS) (Shen et al., 2014). There were 616 significant AS events in *smad1*-cKO PGCs compared to those in sibling PGCs (FDR < 0.05 and inclusion level difference > 10%), including 516/616 exon skipping (ES), 19/616 intron retention (IR), 6/616 mutually exclusive exons (MXE), 32/616 alternative 5′ splice sites (A5SS) and 43/616 alternative 3′ splice sites (A3SS) (Fig. 7A; Supplementary Table 2). Then, we identified 512 protein-coding genes displaying differentially spliced transcripts (AS genes for short) according to these 616 splicing events. Furthermore, we identified 660 AS genes in *smad1*-KD PGCs compared with those in control PGCs (Supplementary Table 3). Remarkably, we observed a highly significant overlap (hypergeometric test, *P* = 2.44e-81) between the AS genes in *smad1*-cKO (121/512) and *smad1*-KD (121/660) PGCs (Fig. 7B), such as *ccdc187 (coiled-coil domain containing 187)* that is involved in microtubule anchoring and located in the centrosome (Priyanka and Yenugu, 2021), *mak* (*male germ cell associated kinase*) that encodes a serine/threonine protein kinase related to cell cycle regulation (Wang and Kung, 2012), *tinf2* (*TERF1-interacting nuclear factor 2*0) that encodes one subunit of shelterin complex involved in distinguishing between telomeres and DNA damage (Lange, 2005), among others (Supplementary Fig. 4A). Altered splicing of these genes was confirmed by RT-PCR (Fig. 7C; Supplementary Fig. 4B). Moreover, GO analysis of all these 1,051 AS genes revealed that the terms chromatin regulator, centrosome, DNA damage checkpoint signaling, and cell cycle were significantly enriched (Fig. 7B). Thus, these data hint that loss of Smad1 in PGCs results in altered splicing of genes related to chromatin structure, DNA damage response, and cell cycle.

**Fig. 7.**
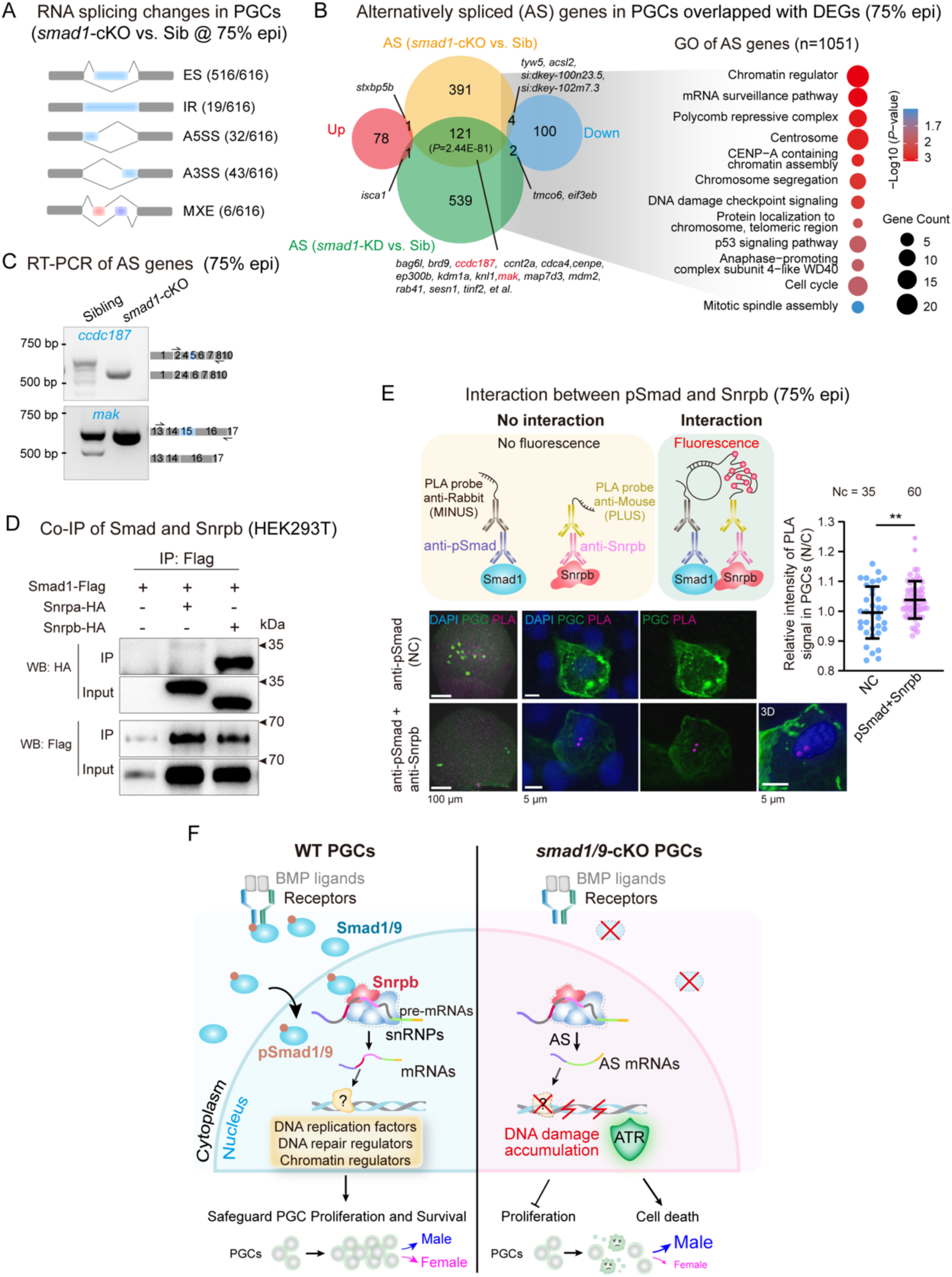
Genes involved in DNA repair, chromatin organization, and cell division show altered splicing patterns in *smad1*-cKO PGCs. (**A**) Schematic summarizing changes in splicing patterns identified by rMATS between control and *smad1*-cKO PGCs at 75% epiboly stage. Changed events were filtered for FDR < 0.05 and absolute percent spliced in (ΔPSI) > 0.1. ES, exon skipping; IR, intron retention; A5SS, alternative 5′ splice site; A3SS, alternative 3′ splice site; MXE, mutually exclusive exon. The proportions of each event type among all 616 significant alternatively spliced (AS) events were shown in parentheses. (**B**) Relatedness of AS genes to DEGs in PGCs. Left, Venn diagrams showing overlaps. Two genes written in red corresponded to those shown in (**C**), with detailed splicing changes. Right, top Biological Process GO terms enriched among alternatively spliced genes. (**C**) RT-PCR validation of altered splicing of selected AS genes in *smad1*-cKO PGCs at 75% epiboly stage. The compositions of exons for the major products were indicated on the right, with the blue exon being alternatively spliced. (**D**) Co-immunoprecipitation (co-IP) of Smad1 with spliceosome components Snrpa and Snrpb. Constructs were transfected into HEK293T cells. (**E**) Detection of the interaction between pSmad1/5/9 and Snrpb in 75%-epiboly embryos by proximity ligation assay (PLA). Top, schematic of PLA principle. Bottom left, representative images showing interaction (PLA probe signals) between pSmad1/5/9 and Snrpb. PGCs were labeled by *kop*:GFP. Bottom right, quantification of nuclear PLA signals normalized to cytoplasmic levels in PGCs. Each dot represented one PGC. Nc, total number of observed PGCs. Error bars indicated SD. Statistical significance was determined by Student’s *t*-test: **, *P* < 0.01. (**F**) Proposed model of BMP–Smad signaling in zebrafish PGC development. WT PGCs proliferate and survive properly, eventually contributing to sex differentiation into either sex. In contrast, *smad1/9*-cKO PGCs exhibit altered transcript splicing and DNA damage, which result in ATR pathway activation, reduced proliferation, and cell death. During the juvenile stage, these PGC defects bias sex differentiation toward males.

To investigate whether Smad1 directly or indirectly regulates alternative splicing in PGCs, we examined the interaction between pSmad and splicing factors both *in vitro* and *in vivo*. First, via co-immunoprecipitation (co-IP) with Smad1 in HEK293T cells, we checked several small nuclear ribonucleoproteins (snRNPs) in core spliceosome components, such as small nuclear ribonucleoprotein polypeptide A (Snrpa) and small nuclear ribonucleoprotein polypeptide B and B’ (Snrpb) (Rodrigues et al., 2023). Co-IP assay revealed a strong interaction between Smad1 and Snrpb, but only a weak or negligible interaction between Smad1 and Snrpa (Fig. 7D). Then, to validate the interaction, we performed proximity ligation assay (PLA) in zebrafish embryos. In brief, when two proteins are adjacent to each other (typically less than 40 nm), specially designed secondary antibodies with attached DNA oligonucleotides (PLA probes) could be close enough for PLA probe ligation, rolling circle amplification, and bound by fluorescently labelled probe, which creates a bright fluorescent spot at the site of interaction (Fig. 7E, upper) (Söderberg et al., 2006). Embryos treated with only pSmad1/5/9 antibody were set as a negative control. Compared with PGCs in the control group, PLA signals were significantly stronger in PGCs within embryos treated with both pSmad1/5/9 and Snrpb antibodies (Fig. 7E). Therefore, these data make it evident that the pSmads might regulate RNA splicing in the nucleus of PGCs by interacting with spliceosome component.

## Discussion

The BMP–Smad signaling pathway is required for PGC induction in mammals, whereas zebrafish PGCs are specified by maternally provided germ plasm. In this study, we observed active BMP–Smad signaling in zebrafish PGCs and demonstrated that BMP–Smad1/9 signaling is dispensable for PGC fate determination and migration but required for PGC proliferation. We generated PGC-specific knockout of *smad1* and/or *smad9* mutants using our recently established double transgenic strategy (Li et al., 2025) to minimize somatic cell interference with PGC development, such as dorsoventral axis formation mediated by BMP signaling. Based on transcriptome analysis and PGC tracing, we conclude that a significant reduction in PGC number, but not PGC-specific gene expression changes and cell migration defects, is triggered by PGC-specific loss of Smad1/9. Notably, ATR–pChk1 is overactivated in *smad1*-cKO PGCs, along with cell cycle arrest and ectopic cell apoptosis, leading to PGC reduction, which can be partially rescued by inhibition of ATR–pChk1. Meanwhile, alternative splicing analysis revealed that Smad1 might regulate mRNA splicing of genes involved in chromatin regulation, cell cycle checkpoint, and DNA damage response, which is validated by direct interaction between Smad1 or pSmad1/5/9 and Snrpb (Fig. 7F). Therefore, we propose that BMP–Smad signaling is fundamental for PGC development across vertebrates but fulfills distinct roles in zebrafish.

In mice, cells in the proximal tier of the epiblast around the gastrulation stage will be induced to commit to the PGC fate by BMP signals from the extra-embryonic ectoderm (Lawson et al., 1999; Matzuk and Burns, 2012; Cantú and Laird, 2013). In contrast, in zebrafish, four pre-PGC clusters (PGC precursors), which inherit sufficient amounts of maternally provided germ plasm, are located separately in early blastula (1k-cell stage), two in the ventral region and two in the dorsal region, and their subsequent proliferation results in PGC population expansion (Aguero et al., 2017). Based on our observations, it appears that, in zebrafish, *smad1/5/9* is dispensable for fate specification and migration of PGCs, and *smad1/9* but not *smad5* are required for proliferation and survival of PGCs. In *smad1/9* KD embryos or PGC-specific cKO embryos, not all PGCs failed to proliferate and died eventually. This could be ascribed to technical limitations of our study. That is, after KD or PGC-specific cKO, the target mRNAs or genes in some PGCs may escape from translation block or gene editing, allowing these PGCs to proliferate normally. We speculate that no PGCs with biallelic null mutations in *smad1/9* would survive, that is, Smad1/9 in PGCs would be essential for PGC survival. Further improvements in PGC-specific knockout strategies are warranted to achieve complete ablation of Smad1/9 and thus enable a more definitive validation of our hypothesis.

Preserving the high degree of genome integrity and stability in germ cells is crucial for reproduction and species continuity. However, PGCs encounter a high risk of DNA damage due to the following three reasons. First, zebrafish embryos undergo rapid cell proliferation, whose cell cycles are fast, ranging from 15 min per cycle during cleavage stages to 30 min per cycle during gastrulation stages (Kimmel et al., 1995). Second, the zebrafish genome is comparable in size to those of humans and mouse (Howe et al., 2013), resulting in extremely high DNA replication stress. Third, zebrafish PGCs also initiate their zygotic genome activation before migration and then go through extensive transcriptome and epigenetic reprogramming during migration to accomplish the lineage specification process (D’Orazio et al., 2021). The combination of these factors poses a significant risk of DNA damage, which increases the need for robust DNA repair in zebrafish PGCs. One interesting question is whether BMP signaling-mediated genome integrity maintenance is germ cell-specific or ubiquitous in early embryos. Notably, we also observed cell death or reduced body length in embryos injected with low doses of either triple *smad1/5/9* MOs or individual MOs, without significant dorsoventral axis defects (Fig. 2B–C; Supplementary Fig. 1B). Recently, it has been reported that BMP signaling relieves DNA replication stress promoting cardiomyocyte proliferation during heart regeneration in zebrafish (Vasudevarao et al., 2025). This ability of BMP signaling to overcome replication stress might also be conserved in mammalian cells (de Jaime-Soguero et al., 2024; Vasudevarao et al., 2025).

Previous studies on how the BMP signaling regulates the cell cycle or DNA damage repair have typically focused on transcriptional regulation by pSmad1/5/9 (Pardali et al., 2005; Su et al., 2009; Kaneda et al., 2011; Vattulainen-Collanus et al., 2018; Lyu et al., 2025). In our study, we observed PGC reductions during the epiboly and somite stages, while no obvious transcriptome changes were detected at 75% epiboly stage, hinting that BMP–Smads signaling might regulate PGC proliferation and DNA damage response through mechanisms other than classical transcriptional regulation. Intriguingly, we detected abnormal splicing events in both *smad1*-cKO and *smad1*-KD PGCs. Meanwhile, it has been shown that Smad3 regulates alternative splicing by directly associating with hnRNPE1, promoting epithelial-to-mesenchymal transition and metastasis in mouse (Tripathi et al., 2016). Thus, we tested whether Smad1 could directly interact with splicing factors and found that it binds Snrpb. However, more detailed investigations are warranted to reveal how BMP–Smad regulates alternative splicing in zebrafish PGCs in the future. Altogether, our findings, along with the systematic PGC-specific *smad1/5/9* knockout and analysis, may help pave the way for future studies of BMP–Smad signaling pathway in PGC development.

## Methods

### Zebrafish husbandry and strains

All zebrafish husbandry and experimental procedures were approved by the Tsinghua University Animal Care and Use Committee. Wild-type Tübingen (TU) strain was used for this study and for generating transgenic and mutant lines. Adult zebrafish were maintained in a water-circulating system at 28.5°C under a 14-h light/10-h dark cycle. Embryos were raised at 28.5°C in Holtfreter’s solution (0.05 g/L KCl, 0.1 g/L CaCl₂, 0.025 g/L NaHCO₃, and 3.5 g/L NaCl, pH 7.0). For imaging of PGCs, 0.003% phenyl-2-thiourea (PTU; Sigma-Aldrich, catalog No. P7629), a tyrosinase inhibitor, was added to the Holtfreter’s solution from 24 hpf onward to inhibit pigmentation.

### mRNA synthesis

For the *smad1-*cKO rescue experiment, an expression vector containing the *smad1* coding sequence followed by the *nanos3* 3’UTR was constructed. These two elements were amplified from a zebrafish cDNA template and inserted into a pXT7 backbone using the Gibson Assembly method. Capped mRNAs were generated from linearized plasmids using mMessage mMachine T7 or SP6 kit (Ambion, ThermoFisher catalog No. AM1344; AM1340) at 37°C for 2–3 h. DNase I treatment was applied to degrade the template DNA, and the synthesized mRNA was purified using the RNeasy Mini Kit (Qiagen, catalog No. 74104). Purified mRNA was aliquoted, diluted, and stored at -80°C.

### Morpholinos and embryo microinjection

Embryo microinjection was performed at 1-cell stage. Injection materials and quantities used were as follows: 1) transgenesis: 10–30 pg *Tol2*-based vector DNA and 150 pg *transposase* mRNA per embryo. 2) mRNA overexpression and MO injection: dosages were provided in the figure legends. The sequences of the MOs were reported before and as follows: *smad1* MO: 5’-AGGAAAAGAGTGAGGTGACATTCAT-3’ (Dee et al., 2007); *smad5* MO: 5’-AACAGACTAGACATGGAGGTCATAG-3’ (Lele et al., 2001); *smad9* MO: 5’-TGCATCGTGAAACGGGTTGATTTTA-3’ (Min et al., 2021); standard control MO: 5’-CCTCTTACCTCAGTTACAATTTATA-3’.

### Inhibitor treatment

Inhibitors were dissolved in DMSO, and stock solutions were freshly diluted in Holtfreter’s solution before use. Embryos treated with corresponding concentrations of DMSO were used as controls.

To inhibit BMP signaling, embryos were treated with 2 μM LDN-193189 (Selleck, catalog No. S2618) during the indicated developmental stages. To inhibit ATR–pChk1 checkpoint activation, embryos were treated with 2 μM AZ20 (Shanghai Yuanye Bio-Technology, catalog No. S81255) from the sphere stage to either 1 or 2 dpf. To inhibit ATM–pChk2 checkpoint activity, embryos were treated with 2 μM KU-60019 (Selleck, catalog No. S1570) from the sphere stage to 2 dpf.

### Generation of PGC-specific knockout mutants

We employed previously established double-transgenic systems (Li et al., 2025) to achieve PGC-specific knockout of *smad1* or *smad9* in zebrafish. For early-stage PGC analysis, *Tg(piwil1:zCas9-nos3 3’UTR)*;*Tg(kop:Eos-CAAX-nos3 3’UTR)* females were crossed with *Tg(U6:smad gRNAs;gsc:tdTomato)* males. cKO embryos were selected based on tdTomato fluorescence during gastrulation, and Eos fluorescence enabled PGC visualization from the dome stage to approximately 2 dpf. For long-term PGC analysis, *Tg(piwil1:zCas9-nos3 3’UTR)*;*Tg(piwil1:mCherry-CAAX-nos3 3’UTR)* females were crossed with *Tg(U6:smad1 gRNAs;cryaa:CFP)* males. mCherry expression allowed persistent PGC labeling, while CFP fluorescence in the lens (detectable from about 30 hpf) was used to distinguish cKO embryos from siblings.

Females were validated based on PGC-specific fluorescence and *cas9* ISH detected in their progeny. To generate *smad1&*9 double knockouts, *Tg(U6:smad1 gRNAs;gsc:tdTomato)* and *Tg(U6:smad9 gRNAs;cryaa:CFP)* lines were crossed to obtain double gRNA-expressing males. These males were then mated with *Tg(piwil1:zCas9-nos3 3’UTR);Tg(kop:Eos-CAAX-nos3 3’UTR)* females to generate *smad1&*9-cKO embryos, which are selected by double-positive CFP and tdTomato fluorescence.

### Immunostaining in embryos

Embryos (before 1 dpf) were collected and fixed in 4% paraformaldehyde (PFA) overnight at 4°C with gentle shaking. After fixation, they were washed three times in PBS for 15 min each at room temperature (RT), dehydrated through a methanol gradient (30%, 50%, 75%, 100% in PBST (0.2% Triton X-100 in PBS)), and stored in 100% methanol at -80°C for at least 30 min. Rehydration was performed through a reverse methanol gradient. Embryonic chorions were manually removed in PBST, followed by antigen retrieval with 10 mM sodium citrate (pH 6.0) at 95°C for 10 min. After cooling naturally to RT, embryos were washed in PBST and blocked (1% bovine serum albumin (BSA), 10% normal goat serum in PBST) for 1 h at RT. Embryos were incubated with primary antibody diluted in blocking buffer overnight at 4°C. Primary antibodies included rabbit anti-pSmad1/5/9 (1:200, Cell Signaling Technology, catalog No. 9511S), rabbit anti-Ddx4 (1:1,000, GeneTex, catalog No. GTX128306), mouse anti-Ddx4 (1:500), rabbit anti-pChk1 (1:200, Cell Signaling Technology, catalog No. 2348), rabbit anti-pChk2 (1:200, MedChemExpress, catalog No. HY-P80799), rabbit anti-γH2AX (1:200, Cell Signaling Technology, catalog No. 2577S).

For active Caspase-3 detection in 2-dpf embryos, cryo-sectioned slides were prepared. Embryos were fixed, washed, dehydrated in sucrose gradients (15% and 30% sucrose-PBS), and embedded in Optimal Cutting Temperature matrix compound (SAKURA, catalog No. 4583). Embedded embryos were sectioned using a cryostat microtome (Leica CM1900) and air-dried for 30 min. Slides were washed in PBS, followed by antigen retrieval with sodium citrate, blocked in buffer, and incubated with primary antibodies overnight at 4°C. Parafilm was used to prevent evaporation during incubation. Primary antibodies included rabbit anti-active Caspase-3 (1:200, BD Biosciences, catalog No. 559565), mouse anti-mCherry (1:200, EASYBIO, catalog No. BE2026).

After primary antibody incubation, samples (whole-mount embryos or cryo-section slides) were rinsed in PBST three times for 30 min each. Next, samples were incubated with fluorochrome-conjugated secondary antibodies diluted in blocking buffer at 1:200 overnight at 4°C. DAPI staining (1 μg/mL, ThermoFisher) was performed for 10 min. Finally, samples were washed and mounted for imaging using a confocal microscope (Leica TCS SP8 STED 3X or Olympus FV3000).

The secondary antibodies used included Alexa Fluor 647 AffiniPure Goat anti-Rabbit IgG (H+L), Rhodamine (TRITC) AffiniPure Goat anti-Rabbit IgG (H+L), Alexa Fluor 488 AffiniPure Goat anti-Rabbit IgG (H+L), Alexa Fluor 647 AffiniPure Goat anti-Mouse IgG (H+L), Rhodamine (TRITC) AffiniPure Goat anti-Mouse IgG (H+L), and Alexa Fluor 488 AffiniPure Goat anti-Mouse IgG (H+L) (all from Jackson ImmunoResearch Labs).

### EdU proliferation analysis

PGC proliferation was detected using the Cell-Light EdU Apollo643 In Vitro Kit (RiboBIO, catalog No. C10310-2) according to the manufacturer’s instructions. Briefly, manually dechorionated shield-stage embryos were incubated in Holtfreter’s solution containing 500 μM EdUTP until the bud stage. Embryos were then washed in fresh Holtfreter’s solution and incubated at 28.5°C. At 2-somite stage, embryos were fixed in 4% PFA overnight at 4°C. After fixation, excess aldehyde was quenched with 2 mg/mL glycine solution for 5 min RT, followed by washes in PBS, PBST, and PBS. Embryos were stained with Apollo staining buffer for 30 min in the dark, followed by methanol and PBS washes. PGCs were labeled with Ddx4 immunostaining, and nuclei were counterstained with DAPI. The embryos were mounted for confocal microscopy (Olympus FV3000).

### Proximity ligation assay

*Tg(kop:gfp-CAAX-nos3’UTR)* embryos were collected at 75% epiboly stage and processed in accordance with the instructions of the first-day steps of the immunostaining (see “Immunostaining in embryos”), except that the blocking procedure and antibody dilution were performed using the buffers provided in the Duolink PLA kit (Sigma-Aldrich, catalog No. DUO82007 and DUO82008). Embryos were incubated overnight at 4 °C with mouse anti-Snrpb (1:200, Santa Cruz Biotechnology, catalog No. sc-374009) and rabbit anti-pSmad1/5/9 antibodies. Next day, embryos were washed twice in 1× Wash Buffer A (Sigma-Aldrich, catalog No. DUO82049) at RT for 5 min each, followed by incubation with PLA PLUS and MINUS probes (Sigma-Aldrich) diluted 1:5 in the supplied antibody diluent for 1 h at 37 °C. After two additional washes in Wash Buffer A, ligation was performed by incubating each sample with 40 μL ligation solution (1 μL ligase, 8 μL 5× ligation buffer, and 31 μL ddH₂O) at 37 °C for 30 min in a humidified chamber. Following another two washes with Wash Buffer A, amplification was carried out with 40 μL amplification solution (0.5 μL polymerase, 8 μL 5× amplification buffer, and 31.5 μL ddH₂O) for 100 min at 37 °C in a humidified chamber. Embryos were then washed twice in 1× Wash Buffer B (Sigma-Aldrich, catalog No. DUO82049) at RT for 10 min each, followed by a final wash in 0.01× Wash Buffer B for 1 min. PGCs were labeled with chicken anti-GFP antibody (1:1,000, Abcam, catalog No. ab13970).

### Whole-mount *in situ* hybridization (ISH)

Antisense RNA probes were synthesized using the DIG RNA Labelling Kit (Roche, catalog No. 11175025910) following the manufacturer’s protocol, assessed by gel electrophoresis, purified with the RNeasy Mini Kit (Qiagen), and stored at -80°C.

ISH was performed using standard procedures. Briefly, embryos were fixed in 4% PFA, dehydrated and rehydrated through methanol gradients, and manually dechorionated. Proteinase K (1 μg/mL) was applied for 5 min to embryos prior to 2 dpf and for 10 min to embryos at 2–3 dpf, followed by re-fixation and washes. After prehybridization for 4 h at 60 °C, DIG-labeled probes (1 ng/μL for *ddx4*, 1.5 ng/μL for *cas9*) were hybridized overnight at 60 °C. Embryos were then washed, blocked, and incubated with anti-DIG-AP antibody (1:3,000, Roche, catalog No. 11093274910) overnight at 4 °C. Signals were developed using BM Purple (Roche, catalog No. 11442074001), and embryos were imaged after PBS washes and post-fixation.

### Cell culture, transfection, and co-immunoprecipitation (co-IP)

HEK293T cells were maintained in Dulbecco’s Modified Eagle Medium (DMEM; Gibco) supplemented with 10% fetal bovine serum (FBS; Gibco) and 1% penicillin-streptomycin (Gibco), at 37 °C in a humidified incubator with 5% CO₂.

For transfection, cells were seeded in 6 cm culture dishes such that they reached 40–60% confluency at the time of transfection on the following day. Plasmid transfection was performed using VigoFect transfection reagent (Vigorous Biotechnology, catalog No. T001) according to the manufacturer’s instructions. For co-IP assays, cells were transfected with the following plasmid combinations: (1) 3 μg of zebrafish *smad1* coding sequence fused with a C-terminal 3×Flag tag (*smad1*-3×flag); (2) 3 μg of *smad1*-3×flag and 3 μg of plasmid encoding zebrafish *snrpa* with a C-terminal HA tag (*snrpa*-HA); or (3) 3 μg of *smad1*-3×flag and 3 μg of plasmid encoding zebrafish *snrpb* with a C-terminal HA tag (*snrpb*-HA). After 24 h, the culture medium was replaced with fresh DMEM.

Cells were harvested 48 h after transfection and washed three times with cold PBS. They were lysed on ice for 10 min in 500 μL lysis buffer (25 mM Tris-HCl, pH 7.2; 150 mM NaCl; 5 mM MgCl₂; 1% NP-40; 5% glycerol). After centrifugation at 12,000 × g for 12 min at 4 °C, the supernatant was collected. For input, 10 μL of lysate was mixed with SDS loading buffer, boiled at 100 °C for 5 min, cooled, and stored at 4 °C until further analysis. For immunoprecipitation, pre-washed Pierce Protein A/G Magnetic Agarose Beads (Thermo Fisher Scientific, catalog No. A36797) were added to the remaining lysate (≥5 μL beads per sample) and incubated overnight at 4 °C with gentle rotation. Beads were collected by brief centrifugation (1,000 × g), washed 5–6 times with lysis buffer, and bound proteins were eluted in SDS loading buffer by boiling at 100 °C for 5 min. Samples were spun briefly and cooled on ice.

Proteins were resolved by 12.5% SDS-PAGE (Yeasen, catalog No. 20326ES), transferred to nitrocellulose membranes, and blocked for 1 h at RT with 5% BSA in TBST (50 mM Tris-HCl, pH 7.6; 150 mM NaCl; 0.1% Tween-20). Membranes were incubated with primary antibodies overnight at 4 °C, followed by three washes with TBST (20 min each) and incubation with HRP-conjugated secondary antibodies for 2 h at RT. After thorough washing in TBST, signals were developed using an enhanced chemiluminescence detection kit (Beyotime, catalog No. P0018S).

Primary antibodies included mouse anti-DDDDK (Flag) tag (1:10,000; MBL, catalog No. M185-3L) and mouse anti-HA tag (1:3,000; MBL, catalog No. M180-3). The secondary antibody was HRP-conjugated goat anti-mouse IgG (H&L) (1:10,000; EASYBIO, catalog No. BE0102).

### Fluorescence-activated cell sorting (FACS) assay

Transgenic embryos expressing fluorescent protein-labeled PGCs at desired stages were collected and dechorionated using protease treatment. Embryos were transferred to a 1.5-mL RNase-free Eppendorf tube and washed three times with Holtfreter’s solution, followed by three washes with nuclease-free PBS. For early-stage embryos (before 1 dpf), single-cell suspensions were prepared by gentle pipetting in 500 μL of pre-chilled nuclease-free PBS. For embryos after 1 dpf, a 10-min incubation in 0.25% trypsin at 28°C preceded trituration. Samples were centrifuged at 500 × g for 5 min at 4°C, the supernatant was discarded, and the pellet was resuspended in pre-chilled PBS or PBS with 10% fetal bovine serum to halt trypsin activity. This washing and centrifugation process was repeated 2–3 times until the pellet turned white. Cell suspensions were passed through a 35 μm mesh filter (Falcon, catalog No. 352235) to remove clumps. Cells were stained with DAPI to distinguish live and dead cells, and the suspension was sorted using a MoFlo Astrios EQ Cell Sorter.

### RNA extraction from PGCs

For RNA extraction from FACS-sorted PGCs (approximately 3,000 cells), TRIzol reagent (Invitrogen, catalog No. 15596018) was used. Cells were lysed in 500 μL TRIzol, incubated, and then mixed with 100 μL chloroform. The aqueous phase was collected after centrifugation and mixed with isopropanol and 10 μg RNase-free glycogen (Invitrogen, catalog No. 10814010). Following incubation at -20°C, RNA was pelleted by centrifugation, washed with 70% ethanol, air-dried, and dissolved in nuclease-free water. RNA integrity was assessed by agarose gel electrophoresis, and the concentration was measured. RNA was stored at -80°C.

### Quantitative RT-PCR

Oligo(dT)-primed first-strand cDNA was synthesized from total RNA using M-MLV reverse transcriptase (Promega, catalog No. A2791) following the manufacturer’s instructions. qRT-PCR was conducted with 3–5 ng of cDNA using TransStart Top Green qPCR SuperMix (TransGen Biotech) on a Roche LightCycler 480 II. Reactions were performed in triplicate to reduce error, and gene expression levels were normalized to *β-actin* using the 2−ΔΔCT method (Livak and Schmittgen, 2001). Primer sequences used for RT-PCR were listed in Supplementary Table 4.

### Assessment of genome editing efficiency by targeted amplicon sequencing

Approximately 300 PGCs or somatic cells were isolated by FACS from 15-somite stage *smad1*-cKO and control sibling embryos. Genomic DNA was extracted and subjected to WGA using the Discover-sc Single Cell Kit (Vazyme, catalog No. N601-01) according to the manufacturer’s instructions. The amplified genome DNA was diluted and used as a template for PCR with Phanta Max Super-Fidelity DNA Polymerase (Vazyme, catalog No. P505) to amplify three target regions of *smad1* gRNAs per sample (Primer sequences are listed in Supplementary Table 4). PCR products were purified by gel extraction. For each group, the three PCR products were pooled in equimolar amounts and used for library preparation with the NEBNext Ultra II DNA Library Prep Kit (NEB, catalog No. E7645S). Libraries were sequenced on an Illumina HiSeq-PE150 platform, generating 150-nt paired-end reads by Novogene.

Sequencing reads were analyzed using CRISPResso2 (v2.3.2) in “CRISPRessoPooled” mode (Clement et al., 2019), with default parameters and substitutions excluded. Notably, the parameter --min_paired_end_reads_overlap was set to 0 to retain paired-end reads even if they lacked any overlapping region. This adjustment was critical because the gRNA target sites were deliberately designed near one end of each amplicon rather than centrally, ensuring that either the forward (R1) or reverse (R2) read alone would confidently cover the target site for editing analysis. Indel frequencies at the target sites were quantified to determine genome editing efficiency. gRNA sequences were listed in Supplementary Table 4.

### RNA-seq library preparation and sequencing

Full-length cDNA from fewer than 400 FACS-sorted PGCs was generated using the Single Cell Full Length mRNA-Amplification Kit (Vazyme, catalog No. N712), according to the manufacturer’s manual. Cells were lysed in 1 μL of lysis buffer with RNase inhibitor, stored at -80°C if necessary, and reverse transcribed using an Oligo(dT) VN Primer. Sc Reverse Transcriptase added adapter sequences to the 3’ end, followed by 18 cycles of cDNA amplification. For library preparation, 1 ng of cDNA was processed with the TruePrep DNA Library Prep Kit V2 (Vazyme, catalog No. TD503), where adaptors were inserted via transposase reaction and tagged fragments were amplified. After size selection (approximately 450 bp) and purification using VAHTS DNA Clean Beads (Vazyme, catalog No. N411), libraries were sequenced on the Illumina HiSeq-PE150 platform, generating 150-nt paired-end reads by Novogene.

### RNA-seq data processing

RNA-seq reads were first subjected to quality control and preprocessing using fastp. This step included adapter trimming and removal of low-quality reads to ensure high-quality data for subsequent alignment. The cleaned reads were then aligned to the zebrafish reference genome (danRer11) using STAR (version 2.7.10b) with the following parameters: --twopassMode Basic --quantMode TranscriptomeSAM GeneCounts --outSAMtype BAM SortedByCoordinate --outFilterMultimapNmax 20 --alignSJoverhangMin 8 --alignSJDBoverhangMin 1 --outFilterMismatchNmax 999 --outFilterMismatchNoverReadLmax 0.04 --alignIntronMin 20 --alignIntronMax 1000000 --alignMatesGapMax 1000000 --outFilterType BySJout. Aligned reads were then counted for each gene using featureCounts (version 2.0.3). StringTie (version 2.2.1) was used to calculate TPM (Transcripts Per Million) values with the following parameters: stringtie -e -B -G Danio_rerio.GRCz11.108.gtf.

To identify differentially expressed genes, read counts were imported into DESeq2 with significance thresholds *P* value < 0.01 and |log2FoldChange| >1. Volcano plots were generated using ggplot2 (version 3.5.1) to display log2(FoldChange) vs. -log10(*P* value).

Functional annotation was conducted using the Database for Annotation, Visualization and Integrated Discovery bioinformatics resource (Huang et al., 2009; Sherman et al., 2022). GO terms for each functional cluster were summarized as a representative term, and the statistical significance of these terms was visualized using *P* values.

### Alternative splicing analysis and validation by PCR

Alternative splicing events were analyzed using rMATS (version 4.1.2) across five basic splicing patterns using two (control vs. *smad1*-KD) or three (control vs. *smad1*-cKO) biological replicates. BAM files were aligned to the zebrafish genome (GRCz11.108.gtf) with the following parameters: rmats.py --b1 control.txt --b2 *smad1*-cKO (or *smad1*-KD).txt --gtf Danio_rerio.GRCz11.108.gtf -t paired --readLength 150. rMATS (version 4.1.2) detects differential splicing by employing a hierarchical framework to model exon inclusion levels (denoted as PSI), accounting for estimation uncertainty in individual replicates and variability among replicates. Differential splicing events were filtered based on FDR ≤ 0.01 and |ΔPSI| > 0.1. Results were visualized using rmats2sashimiplot (version 3.0.0).

To validate selected alternative splicing events, full-length cDNA prepared from FACS-sorted control and *smad1*-cKO PGCs (see “RNA-seq library preparation and sequencing”) was used as template for PCR amplification using Phanta Max Super-Fidelity DNA Polymerase. Amplified products were resolved on 2% agarose gels or subjected to Sanger sequencing. Primer sequences used for validation were listed in Supplementary Table 4.

### Live imaging of embryos

For live imaging before 1 dpf, a small hole was made in the chorion using a 1 mL syringe without fully removing it. This approach stabilized the embryo while maintaining protection, as the broken chorion adhered to the embryo, preventing rotational movement. Embryos were placed in custom-carved wells in solidified low-melting-point agarose in a glass-bottom confocal dish and positioned for optimal imaging. To capture dynamic PGC migration and embryo morphological changes, z-stack scanning was used to minimize loss of PGCs between frames. For embryos post-1 dpf, they were manually dechorionated, anesthetized with 0.02% tricaine, and embedded in 1% low-melting-point agarose in Holtfreter’s solution, which was covered with tricaine-supplemented Holtfreter’s solution.

Confocal and live-cell imaging were performed using a Leica TCS SP8 STED 3X or Olympus FV3000 microscope. Image processing was carried out with Imaris (Oxford Instruments) and ImageJ software. Stained tissues and phenotype images were captured in 5% methyl cellulose (Sigma-Aldrich, catalog No. M-6385) using a CCD camera under a Leica M125 C stereoscope.

### Quantification and statistical analysis

Fluorescence intensity measurements were performed using ImageJ (Version 1.54F). For immunostaining, the central Z-plane was selected, and nuclear regions were delineated using the DAPI signal. Cytoplasmic regions were outlined adjacent to the nucleus. Mean fluorescence intensity was calculated as total intensity (IntDen) divided by the area (Area). RNA-seq included at least two biological replicates. Statistical analyses and graphs were generated using GraphPad Prism 8, with error bars representing mean ± SD in column graphs. Box and whisker plots showed all points from minimum to maximum with the median indicated. Statistical details, including sample sizes and methods, are provided in figure legends. All Student’s *t*-tests were unpaired, and *P*-values were two-sided for both Student’s *t*-tests and Fisher’s exact test.

## Author contributions

Conception and design of the research: A. Meng, X. Wu, T. Zheng. Drafting the manuscript: A. Meng, X. Wu, T. Zheng. T. Zheng performed most experiments. Y. Li, W. Zhang, and C. Xing contributed to the generation of *Tg(piwil1:zCas9-nos3 3’UTR)*. G. Li, Z. Wei, J. Li assisted with the co-IP assay. Z. Jiang helped with the PLA experiment and participated in some discussions. R. Shah assisted with zebrafish husbandry.

## Acknowledgments

This work is financially supported by the National Natural Science Foundation of China (#32588201 to A.M.), the National Key Research and Development Program of China (#2023YFA1800300 to X.W.), and the Yunnan Provincial Science and Technology Project at Southwest United Graduate School (#202302AO370011 to A.M.). We thank all members of the Meng lab for their intellectual and technical support. We thank Dr. Boqi Liu for sharing *Tg(kop:gfp-CAAX-nos3’UTR)*. We are grateful to the Cell Biology Facility and the Sharing Core Facility affiliated with the Center of Biomedical Analysis, Tsinghua University, for technical assistance and daily equipment support.

## Data availability

The RNA-seq data of PGCs are available from NCBI GEO under accession code GSE299653.

To review GEO accession GSE299653:

Go to https://www.ncbi.nlm.nih.gov/geo/query/acc.cgi?acc=GSE299653 Enter token enqrocwqjvefdmn into the box.

