## Supplementary figures for "BMP–Smad1/9 signaling is required for PGC proliferation in zebrafish"

#### Supplementary Material file includes:

Supplementary figures S1 to S4

### Supplementary Figures

#### Supplementary Figure 1

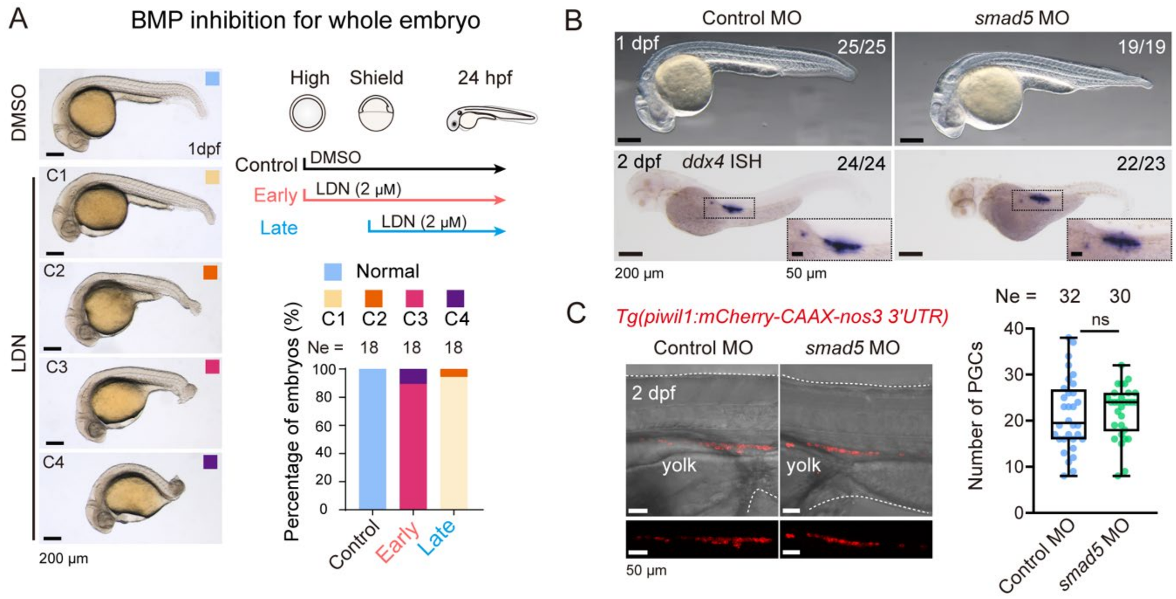


**Supplementary Figure 1: Effects of BMP signaling inhibition and *smad5* knockdown in zebrafish embryos.**

(**A**) Morphological changes caused by BMP inhibitor treatment. Left, categories of body axis defects at 1 dpf. Note variable degrees of posterior trunk malformation. Top right, scheme of treatment with the BMP inhibitor LDN-193189. Bottom right, the ratio of embryos in each category. Ne, number of observed embryos. (**B**) Effect of *smad5* knockdown on *ddx4* expression. Embryos at the 1-cell stage were injected with 0.5 ng control MO or *smad5* MO and observed at 1 dpf (top) or collected at 2 dpf for *ddx4* expression analysis by ISH (bottom). (**C**) Effect of *smad5* knockdown on PGC number. *Tg(piwil1:mCherry-CAAX-nos3 3'UTR)* embryos at the 1-cell stage were injected with 0.5 ng control MO or *smad5* MO. Left, morphology of the gonad region of representative embryos, with PGCs labeled by mCherry. Right, number of PGCs per embryo. Boxes represented the interquartile range with the median indicated by the center line, and whiskers extended from the minimum to maximum values. Each point represented an individual embryo. Ne, number of examined embryos. Statistical significance was determined by Student's *t*-test: ns, not significant ($P \geq 0.05$).

#### Supplementary Figure 2

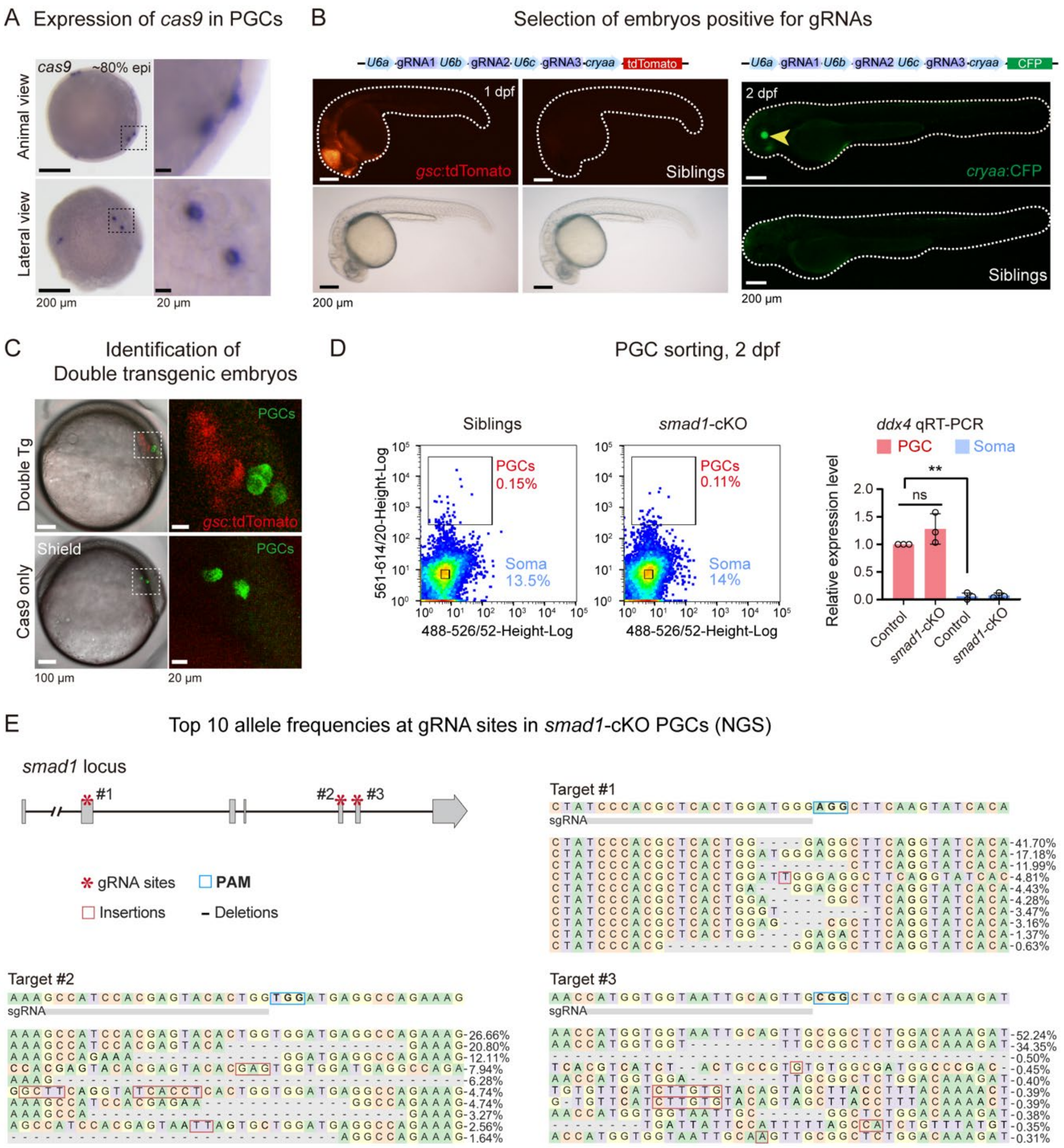


**Supplementary Figure 2: Characterization of the germline-specific knockout system.** (**A**) *cas9* expression in *Tg(piwil1:zCas9-nos3 3'UTR)* embryos at 80% epiboly stage, examined by ISH. Embryos were shown in animal pole (top left) and lateral views (bottom left), and the boxed PGCs-containing regions were enlarged on the right. (**B**) Expression of selection marker in gRNAs transgenic embryos. Two different transgenic lines were shown. Embryos were laterally viewed with anterior to the left. In the right panel, *cryaa*:CFP expression in lens was indicated by an arrowhead. (**C**) Detection of PGCs in double transgenic embryos. Representative images showed *kop:Eos* and *gsc:tdTomato* signals in shield-stage control and *smad1*-cKO embryos, derived from crosses between *Tg(piwil1:zCas9-nos3 3'UTR;kop:Eos-*

*CAAX-nos3 3'UTR)* females and *Tg(U6:smad1 gRNAs;gsc:tdTomato)* males. PGCs were labeled by *kop*:Eos (green). (**D**) FACS sorting and *ddx4* examination of mCherry-positive PGCs and mCherry-negative somatic cells at 2 dpf. Left, representative flow cytometry plots; right, qRT-PCR detection of *ddx4* mRNA levels in sorted PGCs and somatic cells (soma). *β-actin* served as an internal control. Error bars represented SD. Statistical significance was determined using Student's *t* test: ns ($P \geq 0.05$); **, $P < 0.01$. (**E**) Allele frequencies at *smad1* target sites in PGCs from *smad1*-cKO embryos at the 15-somite stage. Data correspond to Fig. 3B.

Supplementary Figure 3

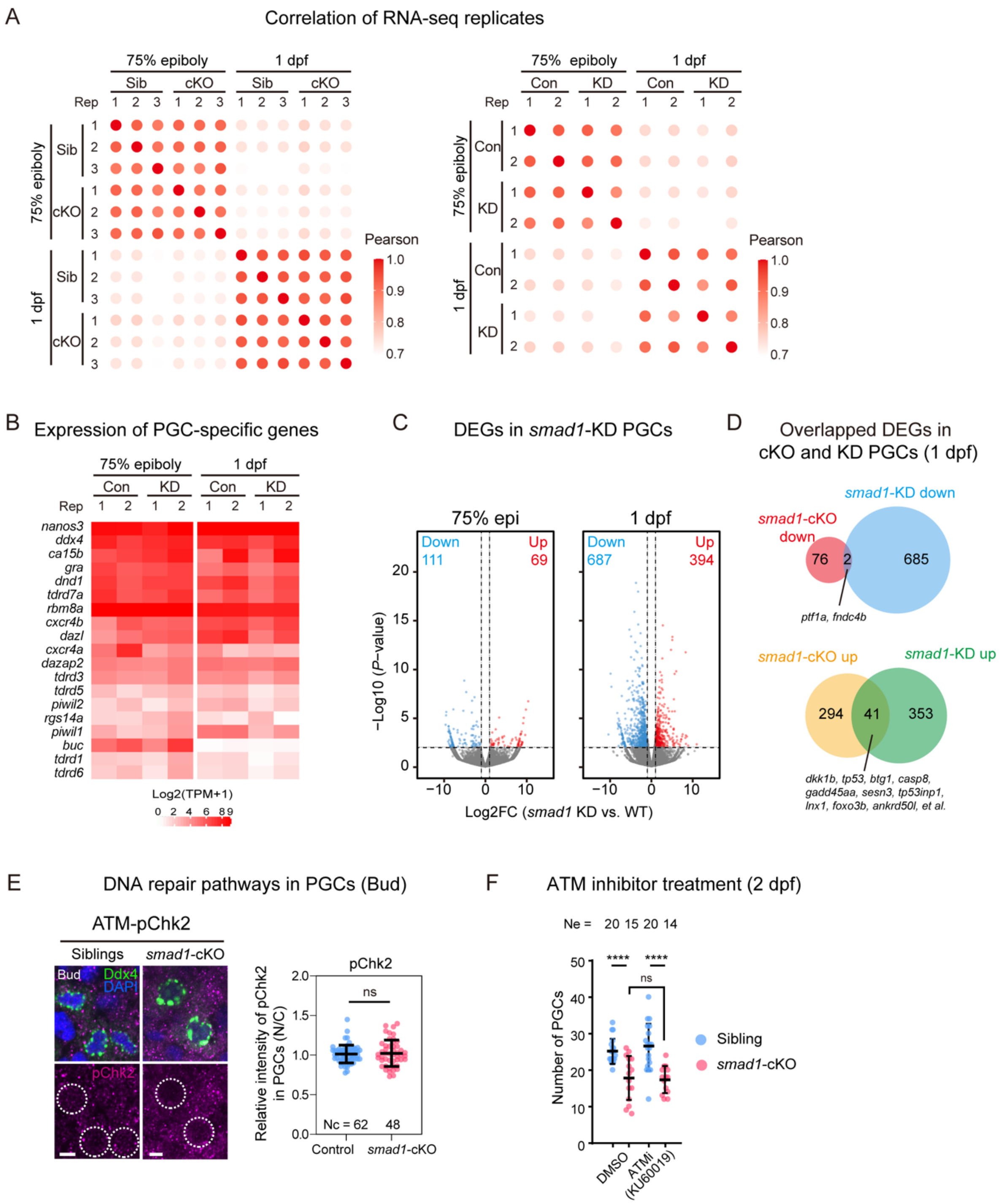


**Supplementary Figure 3: Transcriptome analysis of *smad1*-cKO and *smad1*-KD PGCs and ATM pathway-related phenotypes.**

(**A**) Bubble plots showing *Pearson*'s correlations of RNA-seq data from *smad1*-cKO versus sibling PGCs (left) and *smad1*-KD versus control PGCs (right). (**B**) Expression levels of germline marker genes in *smad1*-KD and control PGCs at 75% epiboly and 1 dpf. (**C**) Volcano plots displaying DEGs in *smad1*-KD PGCs versus control PGCs at 75% epiboly and 1 dpf. Red and blue dots indicated significantly upregulated and downregulated genes, respectively. (**D**)

Venn diagrams showing overlaps between DEGs in *smad1*-cKO versus sibling PGCs and *smad1*-KD versus control PGCs at 1 dpf. (**E**) Representative images (left) and quantification of pChk2 immunostaining (right) in *smad1*-cKO and control PGCs at the bud stage. Nuclear pChk2 signal intensities were normalized to cytoplasmic levels. (**F**) PGC numbers in *smad1*-cKO and sibling embryos treated with DMSO or ATM inhibitor (KU60019). Each dot represented an individual PGC (**E**) or embryo (**F**). Nc, number of PGCs; Ne, number of embryos. Error bars represented SD. Statistical significance was determined using Student's *t* test: ns, not significant ($P \geq 0.05$); ***, $P < 0.001$.

#### Supplementary Figure 4

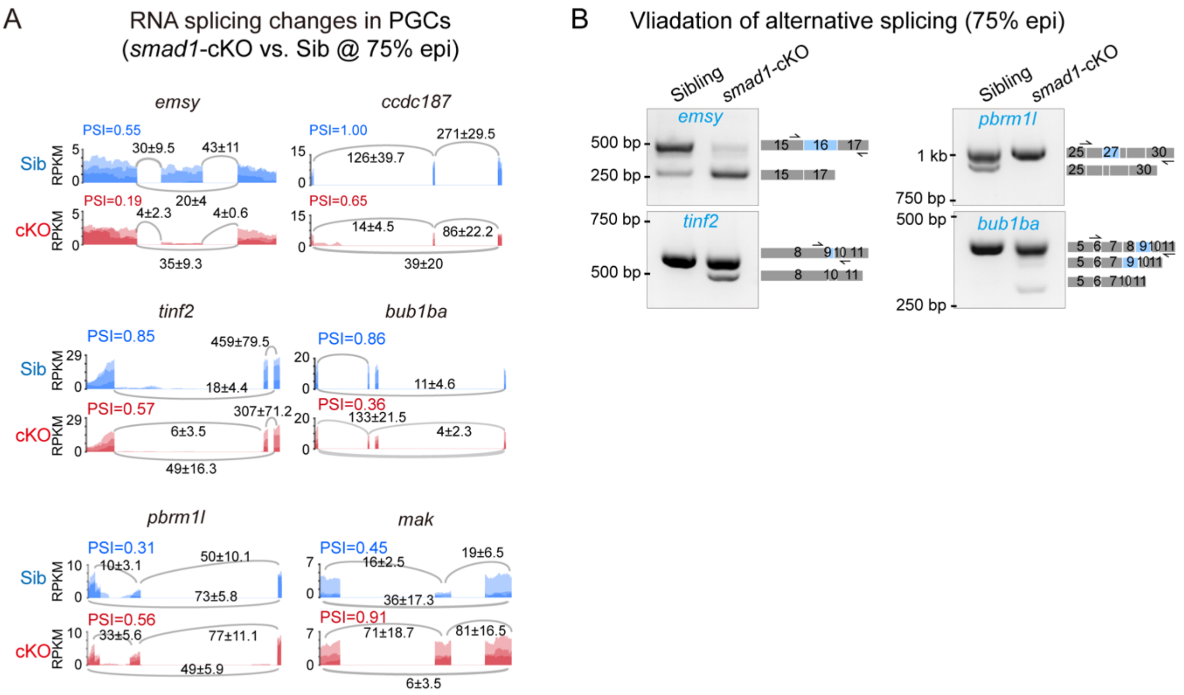


**Supplementary Figure 4: Altered splicing patterns of genes involved in genome maintenance and cell cycle regulation in *smad1*-cKO PGCs.**

(**A**) Sashimi plots showing representative alternative splicing events in *smad1*-cKO versus sibling PGCs at 75% epiboly stage. Numbers indicated junction-spanning read counts (mean ± SD) and percent spliced-in (PSI) values from rMATS analysis across three biological replicates. (**B**) RT-PCR analysis of *emsy, pbrm1l, tinf2,* and *bub1ba* transcripts in sibling and *smad1*-cKO PGCs. Gray and blue boxes denoted constitutive exons and alternatively spliced exons, respectively.
